# Reconstructing signaling histories of single cells via perturbation screens and transfer learning

**DOI:** 10.1101/2025.03.16.643448

**Authors:** Nicholas T. Hutchins, Miram Meziane, Claire Lu, Maisam Mitalipova, David Fischer, Pulin Li

## Abstract

Manipulating the signaling environment is an effective approach to alter cellular states for broad-ranging applications, from engineering tissues to treating diseases. Such manipulation requires knowing the signaling states and histories of the cells *in situ*, for which high-throughput discovery methods are lacking. Here, we present an integrated experimental-computational framework that learns signaling response signatures from a high-throughput *in vitro* perturbation atlas and infers combinatorial signaling activities in *in vivo* cell types with high accuracy and temporal resolution. Specifically, we generated signaling perturbation atlas across diverse cell types/states through multiplexed sequential combinatorial screens on human pluripotent stem cells. Using the atlas to train IRIS, a neural network-based model, and predicting on mouse embryo scRNAseq atlas, we discovered global features of combinatorial signaling code usage over time, identified biologically meaningful heterogeneity of signaling states within each cell type, and reconstructed signaling histories along diverse cell lineages. We further demonstrated that IRIS greatly accelerates the optimization of stem cell differentiation protocols by drastically reducing the combinatorial space that needs to be tested. This framework leads to the revelation that different cell types share robust signal response signatures, and provides a scalable solution for mapping complex signaling interactions *in vivo* to guide targeted interventions.

## INTRODUCTION

Cellular state transition driven by combinations of external signals is the foundation for building tissues in developing embryos, maintaining physiological homeostasis in adults, and triggering disease onset and progression. Identifying which combinatorial signaling conditions trigger state transition of the cells in their endogenous contexts is crucial for understanding and engineering the respective biological process. Genetic perturbations and transgenic signaling reporters are the major ways to correlate signaling pathways with cellular state transitions *in vivo*, but difficult to apply to human tissues ^1,2^. Newer generations of synthetic tools for recording signals have been developed for cells cultured *in vitro*, but their application in animals remains to be tested ^3–5^. Importantly, all the above-mentioned approaches are low-throughput and lacking temporal precision. In many systems, identifying the combinatorial signaling codes that drive cell state transition in time remains a challenging goal. The recent explosion of single-cell RNA sequencing (scRNAseq) and spatial transcriptomics data ushered in numerous computational algorithms to infer cell-cell communication based on gene expression ^6–9^. These methods often rely on the presence of ligand-receptor pairs in the tissue, which does not guarantee signaling pathway activation ^10–12^. Therefore, these methods estimate the signaling potential rather than the signaling state.

Signal activation induces transcriptional responses inside a cell, which is one of the most direct indicators of a cell’s signaling state and compatible for multiplexed high-throughput measurements. However, given the number of signaling pathways, the diversity of cell types, and the combinatorial nature of signaling pathway usage, it is impractical to experimentally measure the transcriptional responses of all cell types upon stimulation by each defined signal. This raises the question of how many signal perturbation measurements are needed before we can accurately infer signaling states of most cell types in their endogenous contexts. While some signaling pathways use common processing machinery across cell types to produce distinct outcomes, the conservation of transcriptional responses in different cell types remains insufficiently explored ^13^. In fact, transcriptional responses to a signaling pathway are often considered unique to each cell type ^14^. If this is true, one would need to perform signal perturbations on the exact cell type of interest, which is often impractical or not scalable for *in vivo* cell types. Therefore, we asked if each signaling pathway has its own transcriptional signature that is shared across diverse cell types. If such signatures exist, it offers a new path for mapping cell signaling states and histories in their endogenous contexts.

Here, we developed a novel experimental-computational framework to learn response signatures from high-throughput *in vitro* perturbations and use these signatures to map the signaling activities in *in vivo* cell types (**Fig. 1a)**. This framework revealed the existence of response signatures shared across diverse cell types, acting as a “fingerprint” for each signaling pathway. Using these “fingerprints”, we mapped the signaling states of over 40 defined cell types in mouse embryos for the first time, revealing patterns of combinatorial code usage with unprecedented temporal resolution and biologically meaningful heterogeneity within each cell type. The framework relies on the design of sequential combinatorial signal screens on human embryonic stem cells (hESC) and modeling the data with a deep neural-network-based algorithm, namely Intracellular Response to Infer Signaling State (IRIS). The inference recovered signaling histories along well-studied cell lineages and made accurate predictions on understudied lineages, such as organ-specific mesenchyme, which was experimentally validated. IRIS greatly reduced the combinatorial signaling space that needs to be experimentally tested to develop hESC differentiation protocols *in vitro*. This new framework represents a conceptual and technical leap in predictive modeling of cell-cell communication that can enable discovery in fundamental biology and design of therapeutic interventions.

**Fig 1.**
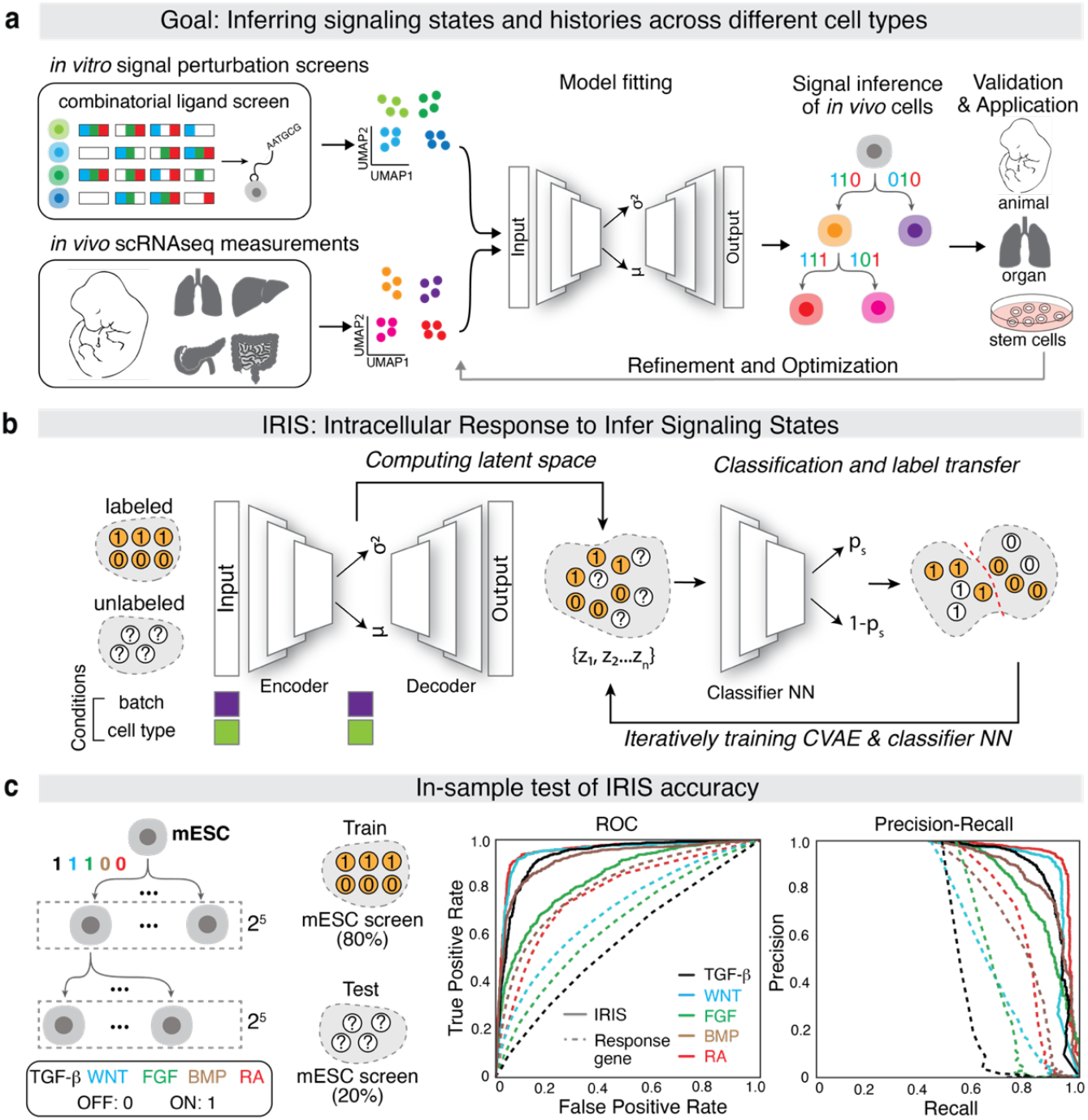
IRIS learns response signatures for accurate signaling state inference. **a**, Schematic of the proposed workflow for inferring signaling histories and optimizing hESC differentiation protocols. *In vitro* high-throughput signal perturbation data sample transcriptional responses to specific signaling pathways, and these response signatures can be learnt by transfer-learning models and used for inferring signaling states among *in vivo* cell types of interest. The inference can iteratively be used to propose new directed stem cell differentiation protocols. **b**, Schematic representation of the IRIS model. IRIS uses a CVAE (conditional variational autoencoder) architecture that includes a feedforward NN (neural network) for classification. This jointly computes a latent space and classifiers for signaling states in the space. **c**, IRIS improves accuracy of signaling state inference over the response-gene method on an 80/20 train-test split of the mESC differentiation data.

### Learning generalizable signal response signatures from the ground up using IRIS

Despite the widely-held belief in the cell-type specificity of signal responses, a small number of genes have been used to indicate signal activation across cell types, such as *Axin2* for Wnt pathway and *Id1* for bone morphogenic protein (BMP) pathway ^15–17^. Therefore, we first asked whether these empirically annotated response genes are sufficient for inferring the pathway activation states across diverse cell types ^17^. We focused on the major developmental signaling pathways, including transforming growth factor beta (TGF-β), fibroblast growth factor (FGF), BMP, Hedgehog (HH), retinoic acid (RA), and Wnt. The activation and inhibition of these pathways occur repeatedly throughout development and in adult tissue regeneration, and thus, they are the major pathways extensively used for pluripotent stem cell differentiation ^18–20^. Using a rare large-scale signal perturbation dataset, in which mouse embryonic stem cells (mESCs) were stimulated with all possible combinations of these six pathways, we created response scores for each pathway in each cell, inferred signaling states, and quantified the inference accuracy (**Extended Data Fig. 1**) ^21^. The response genes had reasonable classification power on RA, WNT, and BMP pathways (>0.75 AUROC), but performed much worse for TGF-β and FGF. The HH pathway was not included because there were too few stimulated cells to render enough statistical power.

When applied to a gastrulating mouse embryo dataset (E6.5-E8.5), the response-gene method recovered some ground-truth knowledge, but in general, it was difficult to convert the response scores into a binary classification (**Extended Data Fig. 2**) ^33^. This is partly because there are no clear cross-cell type heuristics for determining what level of response gene expression constitutes a responding cell. Gene expression measurements are known to be confounded by batch effects, exacerbating the unreliability of the threshold. The small number of response genes used makes them sensitive to the well-known issue of dropouts for scRNAseq data ^22^. Furthermore, these genes were compiled from limited biological contexts that have been extensively studied and may not be generalizable to other cell types. Altogether, these results suggest that shared response signatures might exist across cell types, but such a method based on a small number of genes has limited utility for classifying signaling states.

We hypothesized that this cross-cell type and cross-batch classification problem might be better solved by models of gene expression that are fit to the entire transcriptome as genes do not operate independently of each other ^23^. Models that incorporate the relationship among diverse genes might be more likely to capture the response signatures, be robust to dropouts, and facilitate the choice of thresholds across cell types due to its increased flexibility. Our classification challenge is conceptually similar to the “label propagation” problem in the machine learning field, the goal of which is to learn data features associated with a label and then use those features to annotate unlabeled datasets ^24^. Classic label propagation algorithms have had much success in transferring cell-type labels between different batches of scRNAseq data. In particular, algorithms based on conditional variational autoencoders (CVAEs), such as scANVI and trVAE, can remove the highly nonlinear batch effects more comprehensively and embed cells with similar gene expression patterns together more robustly than comparatively simpler models ^25–27^. They also show success in leveraging small annotated datasets, projecting those onto much larger atlases and learning complicated thresholding rules for diverse cell types.

We asked whether we can apply CVAE-based neural networks to learn generalizable signal response signatures from the ground up. Using the mESC differentiation scRNAseq dataset as a starting point ^21^, we randomly split all the cells into a training set (80%) with signal conditions visible and a test set (20%) with signal conditions hidden. We first applied the CVAE to project all the data across technical batches into a single latent space without awareness of the signal conditions. We then jointly trained the latent space and a classifier network to learn signaling response features, predicted on the test set and compared the prediction with the true signal conditions (**Fig. 1b**). When we fit the model using the default parameters of scANVI, we observed higher classification accuracy than the aforementioned response-gene method (**Fig. 1c**). The model learns a probability of signaling activation for each cell from which it derives a binary prediction simultaneously (**Extended Data Fig. 3**). We named this algorithm IRIS (Intracellular Response Inferred Signaling States).

### Sequential combinatorial signal perturbation screen enables rapid sampling of diverse cell and signaling states

To test the cross-cell-type and cross-dataset performance of IRIS, we generated three orthogonal signal perturbation datasets based on human embryonic stem cell (hESC) differentiation. The mESC dataset produced two main categories of cell states: those resembling primitive streak cells after 24 hrs of exist from pluripotency (mP_d1), and cell types derived from anterior primitive streak (PS) biased towards endodermal fates (mE_d2) ^21^. To maximize cell type/state diversity in the perturbation data, and to produce human cell states that were either related or distinct from those in the mESC screen, we designed a sequential combinatorial signal stimulation strategy spanning distinct developmental stages and lineages (**Fig. 2a**).

**Figure 2.**
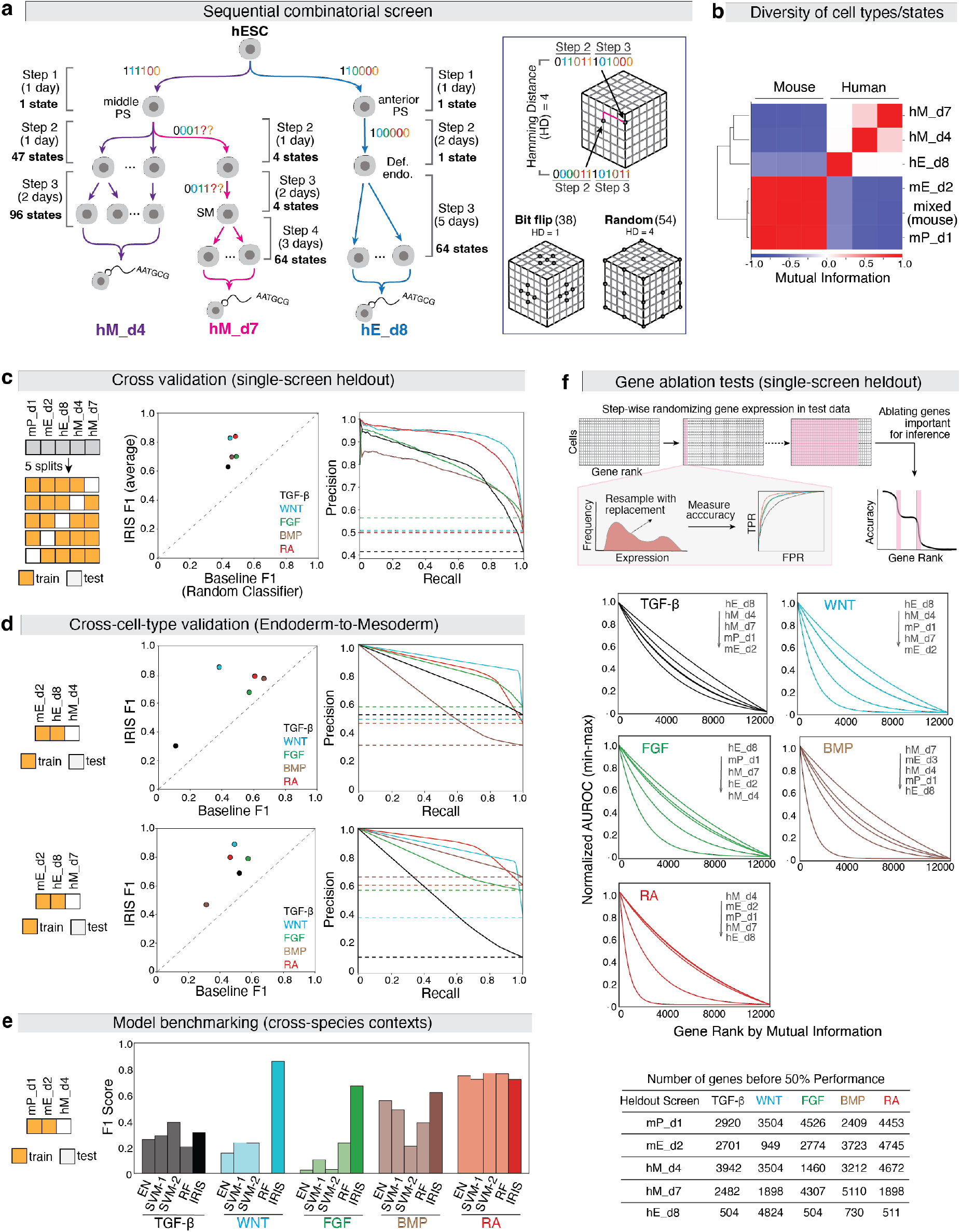
IRIS generalizes across cell types by learning transcriptome-wide signaling signatures. **a**, Design of three sequential combinatorial signaling screens using hESCs. In the last two steps of the hM_d4 screen, signal combinations were represented as points on a 12-dimensional space, with pairwise distances computed using the Hamming distance (HD). Combinations were selected by either a bit-flip strategy (HD = 1 from controls) or a random strategy (HD ≥ 4 apart) (*left*). In the hM_d7 screen, four splanchnic mesoderm populations were generated by varying only RA and SHH at Steps 2 and 3; these RA–SHH pairs were then carried through Step 4, where all 16 possible combinations of TGF-β, Wnt, FGF and BMP were applied to each RA–SHH pair. In hE_d8, all conditions originated from a common Step 2 population and were stimulated with all 64 possible combinations of six pathways in Step 3. hM_d4 and hM_d7, human mesoderm day 4 or 7 respectively; hE_d8, human endoderm day 8; PS, primitive streak; Def endo, definitive endoderm. **b**, Diversity of final cell states in each screen, quantified by transcriptome-wide correlation between distinct screens. **c**, Cross-screen generalization of IRIS when holding out one screen at a time; dots show mean F1 scores across data splits (*left*). Precision– recall (PR) curves show averages across data splits (*solid lines*) compared with a random baseline (*dashed lines*) (*right*). **d**, Cross-cell-type generalization of IRIS trained on endodermal screens (mE_d2, hE_d8) and tested on mesodermal screens (hM_d4, hM_d7). **e**, Benchmarking against other hyperparameter-optimized machine learning models in a cross-species context. EN, Elastic Net; RF, Random Forest. SVM1 and SVM2, two SVM models that differed in hyperparameters. **f**, Gene ablation analysis in the cross-validation test by holding out individual screens. Genes were ranked by mutual information and expression levels were progressively randomized by resampling with replacement. IRIS accuracy was computed after each round of randomization. Results for each data split and each pathway were fit to an exponential function and normalized between 0 and 1 for visualization. The number of genes required to reduce accuracy by 50% reflects the transcriptomic information density. AUROC, area under the receiver operating curve.

Two screens captured mesodermal states after four or seven days of differentiation (hM_d4 and hM_d7, respectively), and one captured endodermal cell states (hE_d8). In both mesodermal screens, hESCs were first induced into middle PS cells that are biased towards mesodermal fate (Step 1), distinct from the anterior PS in the mESC screen ^17,19^. In hM_d4, these cells were exposed to 46 combinations of the six signaling pathways for one day (Step 2), and each condition was further differentiated by a single or multiple signal combination for two days, yielding 96 final states (Step 3). The hM_d7 screen contained cells three days older, in which cells were first differentiated to four splanchnic mesoderm subtypes (Step 1-3) and then subjected to combinatorial stimulation for three days (Step 4). The hE_d8 screen diverged at Step 1, generating anterior PS as in the mESC screen. Those cells were then differentiated into definitive endoderm before exposure to 64 combinatorial conditions over five days. Each final state was barcoded and pooled within each screen for multiplexed scRNA-sequencing ^28^.

Sampling of signal combinations was tailored to each screen. In hM_d4, we implemented a two-step sequential combinatorial design. With 6 signaling pathways acting in two sequential steps, the number of possible combinations is 2^12, which exceeds the current throughput within a single experiment of hESC differentiation. To maximize informativeness, we represented each Step 2-Step 3 sequential combination as a 12-digit binary string and used strategies to spread conditions across the combinatorial space (**Fig. 2a)**. We included 4 published protocols as controls, 38 variants differing by a single ‘bit’ from controls, and 54 random combinations ensuring a minimum Hamming Distance of four between any two conditions ^17^. In hM_d7 screen, BMP, Wnt, TGF-β and FGF specify splanchnic mesoderm, whereas RA and SHH influence the organ identity of the emerging splanchnic mesoderm ^17,29^. Thus, we first generated four splanchnic mesoderm subtypes by varying only RA and SHH, and then applied all 2^4 combinations of the remaining pathways to each subtype in the final step, yielding 64 distinct final states. For hE_d8, we produced a definitive endoderm progenitor state after three days and then sampled the complete 2^6 combinations in the final step. Across all screens, diverse final states were observed (**Extended Data Fig. 4**).

Hierarchical clustering of average cell profiles across screens revealed that human mesodermal datasets (hM_d4, hM_d7) were more similar to each other than to the human endodermal dataset (hE_d8), while cross-species differences exceeded lineage differences (**Fig. 2b**). Human and mouse datasets remained distinct, even between corresponding endodermal states (hE_d8 vs. mE_d2). Together, these screens generated diverse cell states that form a rich ground-truth basis for model evaluation and training.

### IRIS can generalize across batches, cell types and species

With the new perturbation data, we asked if IRIS can truly learn response signatures that generalize across cell types. Given the well-known batch effect between datasets that are collected separately, we first optimized IRIS to minimize the batch effect. Taking advantage of the fact that the mESC data was collected as three independent batches, we performed a hyperparameter search to evaluate how the IRIS architecture affects the cross-batch performance by holding out one batch at a time. Notably, we found that shallow architectures with fewer layers performed the best in this test **(Extended Data Fig. 5)**.

Using optimized hyperparameters, we performed comprehensive cross-validation by withholding one screen at a time. IRIS showed consistently strong performance across all data splits, demonstrating its ability to generalize effectively to unseen datasets (**Fig. 2c, Extended Data Fig. 6)**. Despite pronounced transcriptomic divergence between training and test populations (**Fig. 2b**), IRIS was able to generalize across cell types. To further assess cross-cell-type generalization, we trained IRIS exclusively mesoderm cells are developmentally distinct and non-interconvertible, this represents a stringent test of model generalization. Even under this condition, IRIS substantially outperformed random classifiers (**Fig 2d, Extended Data Fig. 7)**. These results demonstrate that IRIS captures conserved and transferable transcriptional signatures across distinct cell states.

To assess whether IRIS can accurately infer signaling states across species, we trained IRIS exclusively on mouse or human data and tested on the other species. IRIS still successfully captured response signatures generalizable across species (**Extended Data Fig. 8a, b**). Notably, performance improved markedly when a small portion of labeled in-species data from distinct cell types was included during training (**Extended Data Fig. 8c**). Furthermore, we found that including out-of-species data also improved in-species predictions (**Extended Data Fig. 9**). These findings suggest that exposure to diverse cell states, whether within or across species, enhance IRIS’s ability to learn robust, transferable response signatures.

We next compared IRIS with simpler machine learning models optimized using the same hyperparameter tuning procedure (**Extended Data Fig. 10a)**. IRIS performed comparably or modestly better on the in-sample test using mESC data and cross-validation using all screens (**Extended Data Fig. 10b, c)**. However, IRIS maintained stronger performance in more challenging, data-limited cross-species settings (**Fig. 2e)**. Because IRIS more consistently outperformed each of the other models across splits of data and signaling pathways (p < 0.05), we chose IRIS for the subsequent applications.

Finally, we examined the possibility of signal carry-over resulting from prior activation of the same signaling pathway, leveraging the sequential design of hM_d4. Prior stimulation led to an increased false positive rate (FPR) for BMP, WNT and TGF-β pathways, it decreased the FPR for RA and exerted no significant effect on FGF (**Extended Data Fig. 11**). Importantly, even in cells with prior exposure, the FPRs remained below 0.3 for most pathways. These observations indicate that, although signal carry-over may contribute to false-positive classifications, its influence is contingent on cell type.

### Signaling response signatures reflect transcriptome-wide features

The broad generalizability of IRIS prompted us to investigate what constitutes the signaling response signatures. We first asked whether these signatures rely on a small subset of genes and to what extent the previously annotated response genes contribute **(Extended Data Fig. 1a)**. To quantify the importance of individual genes, we performed a gene ablation test. Genes were ranked by mutual information, maximal dispersion, or expression level in the cross-validation test (**Fig. 2c)**. The expression level of each gene was then randomized in the held-out screen to assess its impact on prediction accuracy (**Fig. 2f**). Thousands of genes must be randomized before the predictive power falls to random chance, a pattern consistent across all five screens and three ranking metrics (**Fig. 2f, Extended Data Fig. 12a**). While canonical response genes were ranked highly by mutual information, they explained only a small fraction of the model’s predictive power (**Extended Data Fig. 12b**). These results, while indirect, suggest that IRIS captures the distributed influence of a broad gene regulatory network, rather than relying on a limited set of “context-independent” response genes.

To provide an orthogonal interpretation of the response signatures learnt by IRIS, we combined saliency maps with gene set enrichment analysis (GSEA) ^30^. We first examined whether IRIS simply re-learnt canonical gene sets associated with each signaling pathway. For each pathway, genes were ranked by their contribution to the learnt response signature, quantified as the gradient of the output with respect to each gene at different points of training. We then performed GSEA on the weights associated with each predictor and found the classic signaling pathway response gene sets were enriched or nearly enriched in their respective predictors, suggesting that genes canonically associated with each pathway contribute to the response signature of the corresponding pathway (**Extended Data Fig. 13a**). However, these contributions are not exclusive and many signaling-related gene sets showed high enrichment across multiple pathways (**Extended Data Fig. 13b**). Moreover, IRIS response signatures extended broadly across diverse gene sets, including those beyond signaling-associated annotations, supporting that IRIS captures transcriptome-wide features (**Extended Data Fig. 13c**).

### IRIS reconstructs spatiotemporal signaling dynamics of cell fate lineages

Based on the above observations, we trained IRIS on all screens to infer the activities of all five signaling pathways and the 32 possible combinations across ~40 annotated cell-type clusters from gastrulating mouse embryos (E6.5-E8.5) (**Fig. 3a, b)** ^33^. This analysis revealed highly dynamic shifts in combinatorial signaling codes during this developmental window. Notably, a global transition occurred around E7.0 when early-stage combinations disappeared and were replaced by new combinations that persisted through E8.5. WNT, FGF and BMP were active across most time points and annotated clusters, but only in subsets of cells in those clusters. In contrast, TGF-β was statistically significantly associated with early time points while RA activity emerged later, consistent with prior findings that TGF-β receptor knock-out mice exhibited earlier lethality than RA receptor knockouts **(Extended Data Fig. 14)** ^31–33^.

**Figure 3.**
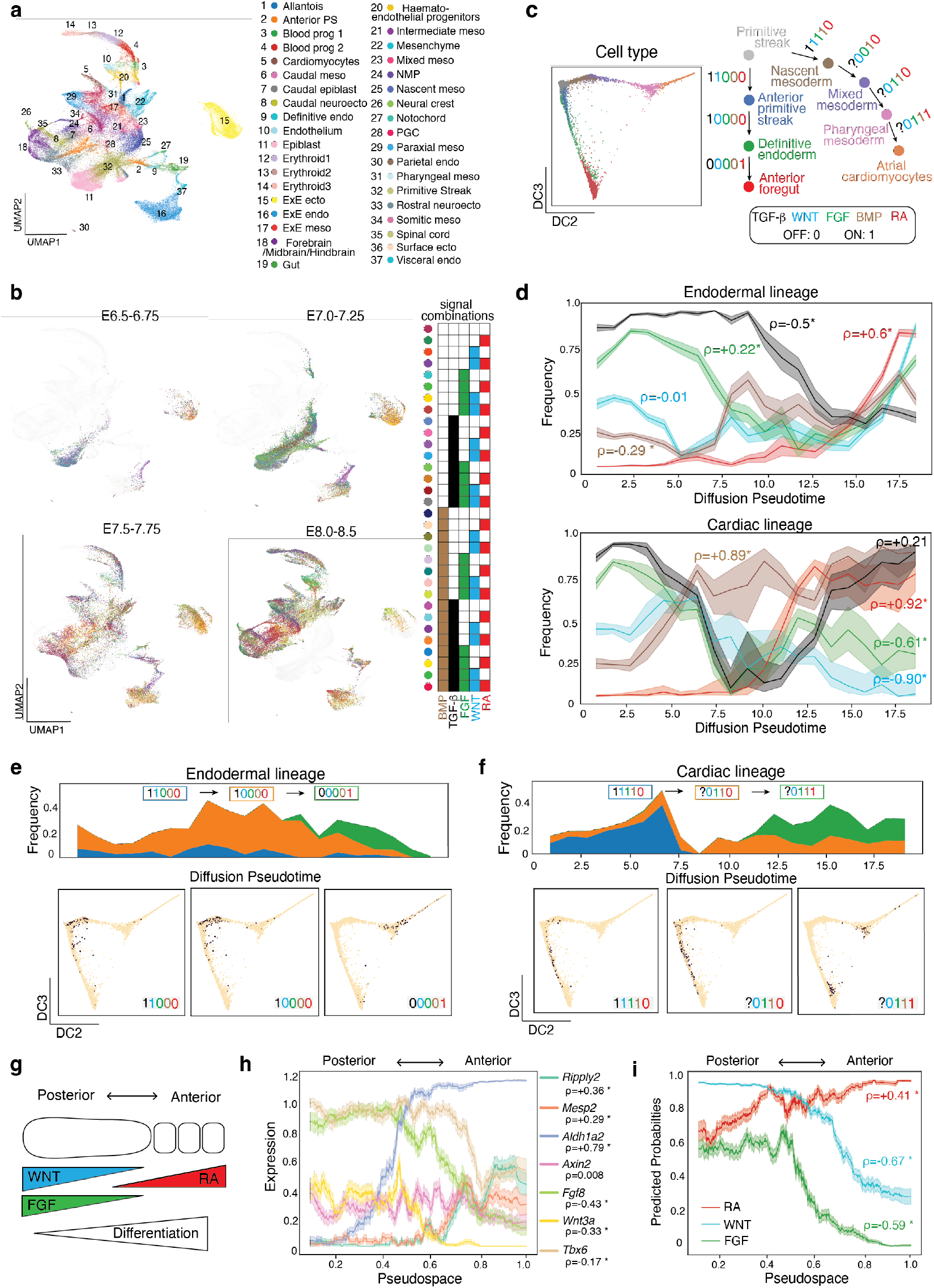
IRIS reconstructs spatiotemporal signaling dynamics of cell-fate lineages. **a**, Annotated cell populations in gastrulating mouse embryos (E6.5-E8.5) ^35^. endo, endoderm; meso, mesoderm; ecto: ectoderm; neuroecto: neuroectoderm. **b**, Dynamic transitions of signal combinations across developmental timepoints. Each combination is color-coded, and cells annotated with that combination are highlighted at each stage. **c**, Diffusion map of endodermal (n=11471) and cardiac lineages (n=18579) (*left*), and corresponding signal combinations during *in vitro* differentiation (*right*). The activation status of TGF-β along the cardiac lineage is unknown. **d**, IRIS-predicted activation frequency for individual signaling pathways along endodermal (*top*) and cardiac lineages (*bottom*). **e, f** IRIS reconstructs the correct temporal order of signal combinations used in differentiation protocols. *Top*, frequency estimates of combinations along each trajectory; *bottom*, diffusion maps showing inferred cells with the corresponding combinations (*black dots*,). Pairwise one-tailed Mann-Whitney U-tests (alternative: greater) on the correct sequence of combinations were significant for both lineages (p < 0.05). **g**, Schematic of three morphogen gradients that spatially instruct somitic-mesoderm differentiation. Progenitors emerge posteriorly and differentiate as they move anteriorly, coupling spatial position with developmental time. **h**, Somitic-mesoderm marker gene expression along pseudospace, approximated by diffusion pseudotime. **i**, Predicted signaling-response probabilities along the inferred pseudospace. Plots showing mean±SEM in **d, h, i**., with * denoting significantly increasing or decreasing trends along temporal or spatial axes, assessed by Spearman’s correlation (p < 0.05).

We next asked whether IRIS could reconstruct signaling histories along diverse cell fate lineages, reflected by sequential usage of signal combinations. The anterior foregut and cardiomyocyte lineages have relatively well annotated signaling histories and are well covered in the mouse gastrulation dataset **(Fig. 3c, Extended Data Fig. 2)** ^34,35^. For each lineage, we selected relevant populations, inferred the activity of each pathway independently, and quantified activation probabilities along differentiation trajectories **(Extended Data Fig. 15a)**. IRIS correctly recovered known temporal trends, including activation of BMP along the cardiac but not the endodermal lineage, inactivation of Wnt preceding cardiomyocyte differentiation, and the activation of RA in both the later endoderm and cardiac populations that correspond to anterior foregut cells and atrial cardiomyocytes respectively **(Fig. 3d, Extended Data Fig. 16)** ^18,19,36^. When comparing inferred signal combinations across pseudotime, their temporal ordering significantly matched expected lineage progression via a permutation test (p < 0.05) **(Fig. 3e, f, Extended Data Fig. 15b)**. For example, the signal combination associated with nascent mesoderm preceded that associated with cardiac specification. These results demonstrate that IRIS can correctly infer signal dynamics and order signal combinations along developmental trajectories.

Unlike other cell-cell communication inference algorithms which use pseudo-bulk analysis, IRIS predicts on individual cells, which allows IRIS to resolve diverse, biologically meaningful cell states within heterogeneous clusters. Within the cardiomyocyte population, IRIS identified enrichment of atrial markers in RA+ cells relative to RA-cells (*Nr2f2, Kcna5*; p < 0.05), consistent with RA’s role in atrial specification (**Extended Data Figure 16b**) ^36^. In endoderm, FGF+WNT+BMP+ cells were enriched for mid-hindgut markers (*Cdx4, Cdx1, Msx1, Msx2, Lef1*; p < 0.05), whereas FGF-WNT-BMP-cells expressed foregut markers (*Sox2, Tbx1*; p < 0.05), consistent with posterior FGF4, BMP2 and Wnt ligand activities promoting hindgut and repressing foregut fate in E7.5-E8.5 mouse embryos (**Extended Data Figure 16d**) ^37^. These results underscore the ability of IRIS to disentangle complex cellular heterogeneity and uncover meaningful biological insights.

Next, we asked if IRIS can identify signaling information associated with cells’ location within a tissue. Somitogenesis is a classic example in which along the posterior-to-anterior axis of the tissue, cells become more differentiated. Within this tissue, three morphogens form concentration gradients that provide the positional information to the cells, with RA coming from the anterior and WNT/FGF from the posterior (**Fig. 3g**) ^38^. To test if IRIS can recover these morphogen gradients, we selected the annotated somitic mesoderm population from the mouse gastrulation dataset and arranged cells along an inferred pseudospatial axis, which accurately recovered the anterior-posterior gene expression patterns, with markers for differentiating somites (*Mesp2, Ripply2*) declined posteriorly and markers for progenitor states (*Tbx6*) declined anteriorly (p < 0.05) (**Fig. 3h, Extended Data Fig. 17a**)^38–40^. Given the spatially graded concentrations of these morphogens, we asked if IRIS can quantitatively predict signaling strength. Instead of using the binary classification, we examined the predicted probability of signaling response, and observed that the averaged probabilities along the spatial axis form gradients analogous to the concentration gradients in the natural tissue (**Fig 3i, Extended Data Fig. 17b**). This result suggests that even though IRIS is trained on binary signaling labels, they may indirectly capture information about signaling strength that correlates with the probability of the prediction.

We further examined the predicted signal combinations at the single-cell level and asked if we can use this combinatorial signaling information to correctly map the location of cells within the tissue. Five combinations, out of the eight possible ones, described most of the cells, with cells in the small RA-Wnt+FGF-population (4% of the cells) likely representing FGF or RA false negatives (**Extended Data Fig. 17c-f**). Placing these populations along the anterior-posterior axis based on the three morphogen gradients recovered the spatial patterns of cell-fate gene expression, demonstrating the ability of IRIS to correctly order signal combinations in space in *in vivo* atlases (**Extended Data Fig. 17g**).

IRIS performed robustly on *in vivo* embryonic endoderm and mesoderm cells, prompting us to assess whether its predictive capacity extends to more distantly related cell types. We first applied IRIS to neuroectodermal cells from the mouse gastrulation dataset, which were not included in the training data. The resulting predictions closely mirrored established models of early neural differentiation (**Extended Data Fig. 18**) ^41,42^. To further evaluate generalizability, we retrained IRIS to predict on a published dataset of organotypic cultures derived from primary adult human bronchial epithelial cells stimulated with diverse signaling cues, including developmental signals ^43^. These cultures comprised multiple cell types with variable competence to respond. When response competence was approximated by the expression of the corresponding signaling receptors, IRIS-predicted positive responders were significantly enriched among receptor-expressing populations (**Extended Data Fig. 19**). Together, these findings demonstrate that IRIS generalizes to cell types outside its training domain.

### IRIS helps optimize hESC differentiation protocol for organ-specific mesenchyme

Finally, we asked whether IRIS can help reveal signaling histories along less-studied cell lineages and inform directed hESC/hiPSC differentiation. Organ-specific mesenchymal cells are a family of enigmatic cell types that are essential for organ development, such as instructing the branching morphogenesis in the lung ^44^. Their developmental histories are poorly understood and thus, efforts in deriving them from hESC/hiPSC to build more realistic organoids have received limited success. Taking advantage of a published dataset on mesenchymal diversity in early mouse organogenesis (E8.5-E9.5), we used IRIS to predict signaling activity during the diversification of organ-specific mesenchyme in the foregut ^17,29^. We found that WNT and BMP are significantly enriched in respiratory mesenchyme compared to all other mesenchymal fates (**Fig. 4a, Extended Data Fig. 20**). Although the role of canonical WNT signaling has been studied in the specification of the lung endoderm and differentiation of lung mesenchymal progenitors into various stromal cell types ^45,46,47^, its role in the initial specification of the respiratory mesenchymal identity was not reported. To test the prediction made by IRIS, we cultured the E9.0 mouse foregut *ex vivo* with either a WNT agonist or antagonist for 24 hrs. WNT activation spatially expanded the expression of key respiratory mesenchymal markers (e.g. *Tbx4* and *Foxf1*), whereas WNT inhibition completely eliminated *Tbx4* expression (**Fig. 4b)**. The expansion of these markers is restricted to the anatomical region immediately adjacent to the lung, where the mesenchymal cells likely share common progenitors with the respiratory mesenchyme, suggesting WNT might regulate the choice between alternative organ-identity of mesenchymal progenitor cells **(Extended Data Fig. 21a**).

**Figure 4.**
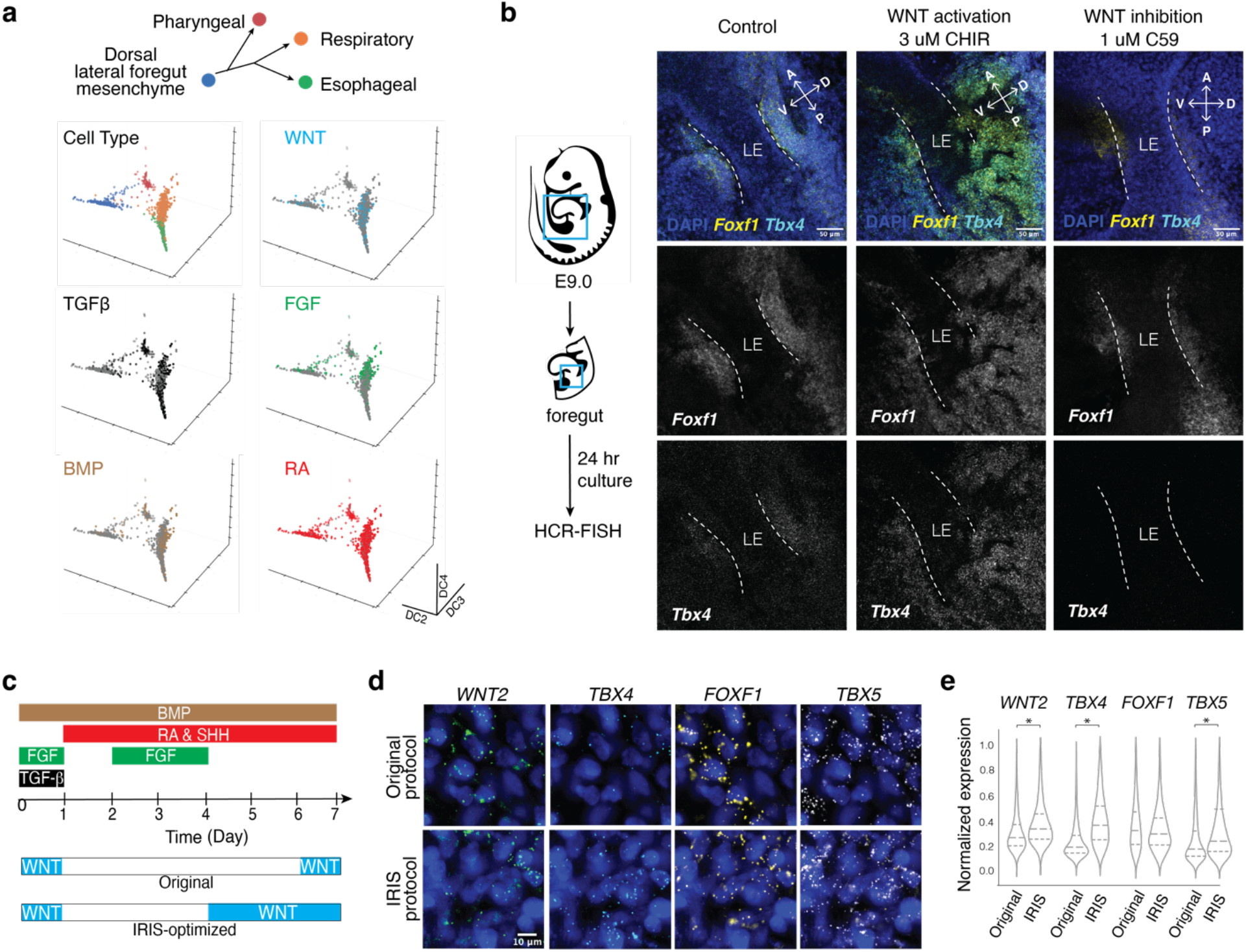
IRIS optimizes hESC differentiation towards organ-specific mesenchyme. **a**, IRIS prediction on early organ-specific mesenchymal cells from E9-E9.5 mouse foregut ^17^. Cells inferred to have the indicated signaling pathway active are highlighted by the corresponding color. WNT and BMP are significantly enriched in respiratory mesenchyme compared to all other mesenchymal fates (Fischer’s exact test, p < 0.05). **b**, Validation of the IRIS prediction using *ex vivo* mouse foregut culture. E9.0 mouse foregut tissues were cultured for 24 hrs with a WNT agonist (CHIR) or an inhibitor (C59), followed by HCR-FISH for respiratory mesenchymal markers in the putative lung region. **c**, Comparison of the original and IRIS-optimized protocols for deriving respiratory mesenchyme from hESCs. The optimized version introduced earlier and extended WNT stimulation while keeping other pathway conditions identical. **d**, HCR-FISH staining of respiratory mesenchymal markers in hESC-derived cells at Day 7 of differentiation (3 µM CHIR in both). **e**, Quantification of **d** showing significantly increased efficiency of respiratory mesenchyme differentiation by IRIS-optimized protocol. * denotes significance by one-sided Student’s t-test (p<0.05)

We sought to apply this knowledge to improve the hESC/hiPSC differentiation protocols for the respiratory mesenchyme, a step towards building lung organoids with branched structure ^29^. The published protocol only applied WNT stimulation at the end of the 7-day differentiation for 24 hrs to promote trachea fate, which occurs after the commitment to respiratory identity (**Fig. 4c**). IRIS predicted that WNT was involved in the commitment to the respiratory fate, a step presumably prior to the trachea fate commitment. We decided to activate WNT earlier and for a longer duration compared to the published protocol, starting from day 4 of *in vitro* differentiation, which corresponds to the stage of splanchnic mesoderm progenitors at roughly E8.5. We found that earlier WNT activation dramatically increases *TBX4* expression in a concentration-dependent manner, and an intermediate WNT activity is optimal for preserving all key marker genes of respiratory mesenchyme, as high Wnt activity decreases *FOXF1* expression (**Extended Data Fig. 21b, c**). The same effect of pro-respiratory fate by early WNT activation holds true across different hESC/hiPSC lines, suggesting that the modified protocol brings us closer to deriving respiratory mesenchyme (**Extended Data Fig. 21d, e**). This shows the utility of IRIS for proposing and optimizing new directed differentiation protocols for stem cells, without having to explore the entire combinatorial space.

## DISCUSSION

Mapping cells’ signaling states and histories is crucial for understanding how cells make fate decisions, but challenging to do due to the expensive and low-throughput nature of such experiments in animal models, and lack of tools in human tissues. We demonstrated a novel experimental-computational framework that performs combinatorial signal perturbation screens on a collection of cell types *in vitro*, and uses the signal response signatures computed at the transcriptome level to infer the activity of the same signaling pathways *in vivo*. The signal response signatures serve as accurate classifiers on diverse cell types, including those not represented in the perturbation screens. This cross-cell-type and cross-batch signal classification is enabled by IRIS, a neural network-based transfer learning algorithm. Although our work focused on six recurring developmental pathways which determine most of cell fate decisions in early embryos, the same framework can be applied to other signaling pathways, such as NOTCH and YAP pathways in developing embryos, cytokines in the immune system, and hormones in regulating physiology.

The fact that signaling response signatures are transferable between unrelated cell types is a surprising discovery, because transcriptional responses to signal activation are often considered unique to each cell type ^48,49^. We demonstrated that the response signatures learnt from mESC and hESC screens can accurately classify signaling states of cells from mouse embryos at equivalent and later stages. To what extent the response signatures learnt by IRIS can be generalized between cells that are far apart in the transcriptomic space remains to be tested. The model itself may be improved by integration with recent advances in transformers and foundational models that are able to learn generalizable principles on large, unlabeled datasets ^50–53^. Increasing cell type diversity in the training data may also enable broader generalization to distant cell types. We envision a cycle of perturbation screens, model training, prediction on new data, and experimental testing, the results of which can be used subsequently to refine the model.

Furthermore, our results support the idea that signaling information can be broadly stored throughout the genome and read out through RNA. Our analysis suggests that many genes carry at least some information about signaling response and classifiers relying on many genes are more robust to cellular contexts and technical noise associated with scRNAseq data. Comparing across pathways, we observed that response signatures to WNT and RA seem to transfer particularly well across cell types in nearly all tests, whereas FGF often, but not always, underperforms. We suspect that this reflects the extent to which a signaling pathway shares its machineries with other pathways. In particular, FGF share signal transduction components with many other receptor tyrosine kinase pathways, whose activities could be present in culture conditions and confound the model. Additionally, our models were fit to data collected after sequential steps and durations of signal stimulation, which might introduce signaling carryover from previous stimulation or an artificial time delay to the predicted signal activation/inhibition respectively. How the response signatures evolve over time remains to be rigorously characterized.

Overall, IRIS is a tool that enables learning signaling states and histories from scRNAseq data. Reconstruction of signaling histories by IRIS relies on accurate lineage inference, which is an active area of algorithm development. Nevertheless, IRIS can act on imperfect cell clusters that include heterogeneous cell types/states, due to its complete reliance on cell-autonomous information at the single-cell level. As a result, IRIS’ response-centric approach is complementary to other signal inference methods relying on ligand-receptor pairing, such as CellPhoneDB, NicheNet and CellChat ^6–8^. These two types of approaches can be used jointly to infer potential cell-cell communication, especially in the case of analyzing spatial transcriptomics data. This will expand opportunities in using scRNAseq and spatial transcriptomics for scientific discoveries and translational applications, such as engineering stem cells and identifying signaling pathways that drive pathological states of the cells.

## Acknowledgements

We would like to thank Qiu Chang Wu, Douglas Lauffenburger, Na Sun, Alexandra Cao, Lauren Tso, Mark Greenwood, and the rest of the Li Lab for helpful discussions. We would also like to thank Jennifer Love, Sumeet Gupta, and Stephan Mraz from the Whitehead Institute Genome Technology core and George Bell from the Bioinformatics and Research Computing core for their advice and assistance throughout the project. This work was supported by National Institute of Health grants DP2HD108777 (P.L.), National Science Foundation Graduate Research Fellowship (N.H.), MathWorks Graduate Fellowship (M.M.), and Allen Family Philanthropies (P.L.).

## Competing interests

Authors declare that they have no competing interests.

## Author contributions

N.H. and P.L. conceived the study. D.F. and P.L. supervised the work. N.H. led the computational and experimental work. C.L. developed the code base. M. Meziane performed experimental validations in mice. M. Mitalipova assisted with stem cell validation experiments.

## Supplementary Materials

### Materials and Methods

#### Single-cell Data Processing

Unless otherwise stated, single-cell data was processed using the Scanpy package (version 1.10.0). For visualization, data was log normalized, scaled according to z-scores, and nearest neighbor graphs were constructed from a PCA computed from these scaled expression values. These nearest neighbor graphs were the basis for the UMAP projections. For developmental lineage visualization, scanpy.tl.diffmap function was used to visualize developmental trajectories. This function takes data, *X* ∈ *R*^*n* × *d*^, and constructs a pairwise similarity matrix between observations using a Gaussian kernel, 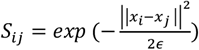. Epsilon is used as the bandwidth parameter. This similarity matrix is then converted to a Markov transition matrix using the partition function 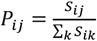. The eigenvectors are then computed from this transition state matrix and used as the diffusion components. Optimal diffusion components were those which, when used in pairs, correctly placed the lineages in order based on known cell type markers and annotations. Human and mouse gene names were converted between each other using mousipy (version 0.1.5). For consistency in the analysis, when mouse and human data was integrated, human gene names were used.

#### Signaling state inference with the response-gene method

Response gene scores were computed by summing the log normalized gene expression values for the response gene sets for each pathway. For a set of j response genes (G) and expression r.

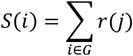

The performance of the inference method was evaluated using the receiver operator characteristic (ROC), precision recall curve (PRC), and F1 score. The ROC was computed and plotted using sklearn.metrics.roc_curve. Specifically, for each signaling pathway, a variety of arbitrary thresholds for the response gene scores were set to classify the pathway activity. Each cell in the test group was classified to be ON or OFF for the pathway activity based on the corresponding response score. By comparing with the ground-truth label, the true positive rate (TPR) and false positive rate (FPR) were calculated for each threshold and plotted against each other to generate the ROC:

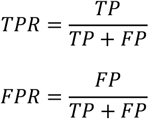

where TP is true positive and FP is false positive. The area under the receiving operating curve (AUROC) was computed using sklearn.metrics.roc_auc_score. The PRC was computed and plotted similarly using sklearn.metrics.prc_curve by setting a variety of arbitrary thresholds for response gene scores. The precision (same as TPR) was plotted against recall at each threshold.

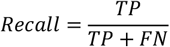

where FN is false negative. The area under the precision recall curve (AUPRC) was computed using sklearn.metrics.auc, which computes the integral over these two scores using the trapezoid rule. F1 score was evaluated using sklearn’s metrics functions (version 1.4.1.post1):

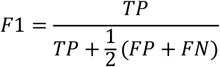

Calculating F1 score requires a single threshold of response gene scores for each signaling pathway, which was computed by finding the threshold which maximally separates true positive (TP) and false positive (FP) according to the ROC curve.

#### IRIS model and hyperparameter selection

IRIS is a semi-supervised model that was implemented using the scANVI architecture^44^ from the scvi-tools package (version 0.20.3), and trained on an NVIDIA-A6000 GPU using the Cuda library (version 11.3). The model architecture is a variant of the conditional variational autoencoder (CVAE) model. It includes an encoder neural network which learns the mapping of the input data X and the conditions S to an n-dimensional latent space Z. A decoder neural network takes as input the conditions S, and latent space encoding in Z. The CVAE model minimizes the evidence lower bound:

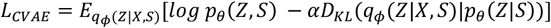

where *q*_*ϕ*_ (*Z*|*X, S*), represents the variational posterior, *p*_*θ*_ (*X*|*Z, S*) is the reconstructed distribution from the latent space, is a hyperparameter which determines the strength of the Kullback-Leibler regularization term (*D*_*KL*_). For the purposes of this analysis, gene expression counts were assumed to follow a zero-inflated negative binomial distribution (ZINB). The default value of 0.01 was used. Simultaneously a feedforward neural network (FNN) takes as input Z and S for each data point and minimizes the cross-entropy (CE) loss:

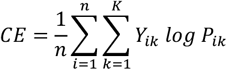

with respect to the target labels Y. Here, k are the individual classes, K is the total number of classes, and P is the probability of being assigned either class label. Each node in the neural network uses a rectified linear activation function *ReLU* = *max* (0, *x*). The final layer of the FNN uses a sigmoid function which computes the probability of the two class labels. These are then thresholded with the value 0.5 to binarize the predictions. The two neural networks are trained jointly to minimize the loss *L* = *L*_*CVAE*_ + *CE*.

Data was stored in the anndata format with observations as individual cells and variables as gene names. *X* ∈ *R*^*n* × *d*^ is the input scRNAseq with raw counts as the values. *S* ∈ *R*^*n* ×*c*^ are the conditions from the covariates c (e.g. batch, cell type, and species), with a one-hot encoding. The target labels *Y* ∈ *R*^1 × *d*^ is presented as a vector in the anndata object with binary indicator variables. X was partitioned into training and test data as specified in each trial. Models were fit by minimizing loss on training data and then evaluated on held-out test data by freezing the model weights and evaluating the prediction output. Model performance was evaluated using the ROC, PRC, and F1 scores computed on the test data (see above).

Hyperparameter selection was performed by fitting distinct architectures on the training data (Layers=1, 2, 3; Hidden nodes=32, 64, 128, 256, 1024; Latent Variables=10, 20, 30, 50, 60). Their performance was evaluated using AUPRC. The best performing architectures were then used in subsequent analysis and evaluated via k-fold cross-validation over the batches. Models were trained for 200 epochs because this was found to provide optimal performance across a range of trials.

#### Setup of the model tests

In-sample tests were performed on random partitions of mESC screen data or the combined mESC and hESC screen data. The data was partitioned using the np.random.choice function to select 20% of the data as test data and the remaining 80% for model fitting. Model performance was evaluated with ROC, PRC, and F1. The cross-validation tests were performed by splitting the mESC screen data into 3 technical batches (mP_d1, mE_d2, mixed) and treating the hESC screen data as three separate technical batches (hM_d4, hM_d7, hE_d8).

In the first test that performs the hyperparameter screen, the model was fit to two of the three mESC batches at a time and tested on the third mESC batch. The test was done iteratively by holding out each batch and averaging the performance metrics across all three held-out batches (**Extended Data Fig. 5, 6**).

The second cross-validation test was performed by including all screen data and iterating through all five possible held-out batches (excluding the mixed batch due to overlap with mP_d1 and mE_d2) (**Extended Data Fig. 7**).

Three tests of out-of-species generalization were performed: (1) The test with limited data was performed by fitting the model with the complete mESC screen data and evaluating on the hM_d screen data, F1 scores were computed and compared to simpler ML model benchmarks. (2) The model was fit to the all of the mESC screen data (mP_d1, mE_d2, mixed) and evaluated on the hESC screen data (hM_d4, hM_d7, hE_d8) and vice-versa, and F1 scores were computed and compared to random baselines. (3) To test if additional target-species data from an unrelated cell type could enable cross-species generalization, the model was fit to mP_d1 plus mE_d2 with or without a random 10% subset of hE_d8 and evaluated on hM_d4. The AUROC and AUPRC scores were compared with and without the subset of hE_d8.

To evaluate if out-of-species data enhanced in-species prediction, two tests were performed. (1) The model was fit to 80% of the mESC screens (mP_d1, mE_d2 and mixed) with and without hM_d4 and evaluated on a held-out 20% partition of the mESC screens. (2) The model was fit to all but a single-held out combination mESC and hESC screen data (mP_d1, mE_d2, mixed, with or without hM_d4) and evaluated on that held out combination. F1 scores were compared with and without hM_d4 in training. To evaluate cross-cell type generalization, the model was fit to the screens representing only endodermal contexts (mE_d2, hE_d8) and evaluated on either hM_d4 or hM_d7. The performance metrics (F1, AUPRC and AUROC) were compared to random baselines.

#### Comparison with classic machine-learning models

Classic machine learning models such as support vector machines (SVM), random forests (RF) and elastic net (EN) were implemented using the sklearn (version 1.4.1.post1) library and run on the same machine. Hyperparameter selection was performed over multiple orders of magnitude for each of the tunable hyperparameters (**Extended Data Fig. 6**).

The support vector machine minimizes the following objective:

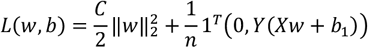

where *w* is the vector of weights, *w* ∈ *R*^*n*^. *N* is the number of data points, 1 ∈ *R*^*d*^ is a vector of ones, *Y* ∈ *R*^*n*^ is a vector of the binary labels, *b* ∈ *R* is the bias term. The scaling factor, C, was chosen as stated in each analysis.

The random forest minimizes the following objective:

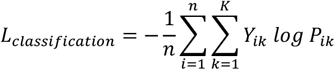

where *K* is the number of classes (in this case for binary classification, 1), *n* is the number of data points, and *P*_*ik*_ is the log probability of the correct classification. The classification loss minimizes the cross-entropy loss.

The elastic net model minimizes the following objective:

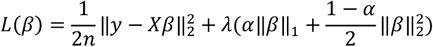

where *β* ∈ *R*^1×*d*^ and are the weights of the model. *N* is the number of data points 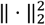 is the euclidean norm and the ‖ ⋅ ‖_1_ is the regularization penalty which is set to values as specified in each trial. α is the weighting parameter for L1 and L2 penalties. For all trials, it was set to 0.5.

#### Computing correlation matrix of transcriptional states and hierarchical clustering

Correlation between cells from two signaling conditions was computed as the averaged gene expression for each gene across both conditions using Pearson’s correlation:

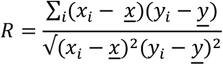

where *x*_*i*_ and *y*_*i*_ are individual genes in the two conditions *x* and *y* respectively. *<u>y</u>* and *<u>x</u>* are the average expression over all genes in the respective conditions. Correlation matrices were then generated with scanpy.pl.correlation_matrix. The distance metric used was 1 − *R*.

All signaling conditions were hierarchically clustered on the dendrogram using Ward’s method. In Ward’s method, two groups are clustered if they result in the smallest increase in total intra-cluster variance, computed as the error sum of squares:

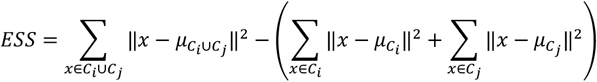

where *C*_*i*_ and *C*_*j*_ represent clusters that are to be merged, and *μ* represents the centroid of a given cluster. This process was continued iteratively until all clusters were merged, representing the root of the dendrogram.

#### Inferring signaling histories along developmental lineages

IRIS was used to infer signaling histories in the mouse gastrulation dataset, foregut mesenchymal dataset, and human airway epithelium ^17, 35^. Cells were first clustered using Leiden clustering in scanpy with default parameters. Cell type annotations were determined using canonical marker genes and annotations provided by each of the datasets. The diffusion components were computed using sc.tl.diffmap and visually inspected for those that placed the cell types and marker gene expression in the expected order consistent with literature. For the cardiac lineage analysis, this included the primitive streak, nascent mesoderm, mixed mesoderm, pharyngeal mesoderm and cardiomyocytes. The endodermal lineage analysis included the primitive streak, anterior primitive streak, definitive endoderm, gut and visceral endoderm. A single diffusion map was made for the endoderm and cardiac lineages. Endodermal cells were sub-clustered and visualized with a diffusion map. The presomitic mesoderm analysis included only cells from the somitic mesoderm cluster and visualized with a diffusion map. The respiratory mesenchyme lineage analysis included only cells from the dorsal lateral foregut, respiratory mesenchyme, esophageal mesenchyme and pharyngeal mesenchyme, which were projected on a single diffusion map.

To infer the signaling histories, two methods were applied. For the response-gene method, response-gene scores were overlaid on diffusion maps to serve as a relative estimate of the pathway activity. Thresholds were chosen to maximize the separation between true and false positives. For the IRIS method, the model was fit to the raw count matrix using optimized hyperparameters for each pathway. Signaling pathway activities were predicted independently for each pathway and the combinatorial signaling state of a cell was composed by concatenating all of the pathway predictions.

To temporally project inferred signaling states along either the developmental trajectories or spatial axes, selected cell populations were lined up across developmental pseudotime, discretized into 35 bins (or smoothed with a bandwidth of 0.1), which was chosen for better visualization. In the case of somitic mesoderm cells, the pseudotime was used as a proxy for a pseudospatial axis from posterior to anterior of the embryo. To validate the pseudotime/psuedospace trajectories, the expression of key cell fate marker genes were projected and analyzed. Finally, the trends of inferred activities among the five signaling pathways were characterized either separately or as combinations. In the case of individual pathways, the binary classification or predicted response probabilities were projected along the pseudo-axis and trends were quantified via Spearman Correlation. The predicted response probabilities are computed by the sigmoid layer of the FNN. In the case of signal combinations, the raw frequency of each inferred combination was plotted across developmental pseudotime, and the temporal ordering of the combinations along these axes were confirmed with Mann-Whitney U-tests of the average position along them. We chose to look at trends and order of signaling combinations rather than abundance as not all progenitor cells go on to become a given final cell fate, making direct interpretation difficult.

#### Feature importance analysis

Feature importance was determined by the reduction of inference accuracy when the expression level of individual genes was sequentially resampled with replacement in the test data. Negative exponential curves of the form *e*^*−αt*^ were fit for visualization using *c*urve_fit in the scipy.optimize library. Resampling with replacement was performed using the numpy.choices function on a vector containing all of the original gene expression values. Genes were ranked from high to low either by their dispersion or by mutual information. Dispersion was computed using scanpy.pp.highly_variable_genes: 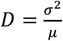, where *σ*^2^ is the variance of the expression for a given gene and is the averaged expression for a given gene. Mutual information was calculated for a given gene X and a given signaling state vector Y, which is a vector of 0 and 1 binary values representing signaling stimulation. The entropy (H) and mutual information (I) were approximated using scikit-learn.mutual_info_classif which implements a k-nearest neighbor estimation of entropy between continuous variables and categorical variables as follows:

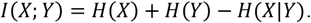

#### Gradients-GSEA Analysis

Gradients were computed natively by Pytorch (version 2.2.2+cu121). Neural networks were trained via backpropagation and the gradient with respect to the input layer was computed as:

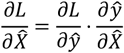

where *L* is the loss function, *y* is the target vector of signaling labels, and *X* is the input data. Genes were ranked according to the magnitude of the gradient. These rankings were used to perform gene set enrichment analysis using GSEApy (version 1.1.3). The normalized enrichment score S is computed as the maximum value of a running sum according to the following equation:

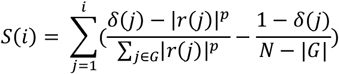

where the score *S* is computed for a given gene set of interest *G, N* is the number of genes in the ranked list and |*G*| is the number of genes in the gene set of interest. The weight parameter *p* indicates how heavily to weigh each gene’s ranking. *δ*(*j*) is the indicator function for whether or not a gene j is in the gene set G. *r*(*j*) is the ranking statistic for a given gene. The normalized enrichment score is then computed as 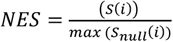. The p value was computed by a permutation test with 1000 samples. The gene set of interest was considered to be enriched if p < 0.05 and nearly enriched if 0.05 < p < 0.06.

#### Stem cell maintenance and directed differentiation

H1 Human embryonic stem cells (hESCs) (WiCells WA01) or induced pluripotent stem cells (iPSCs) (WIBR iPSC66) were maintained in mTESR plus (STEMCELL Technologies 100-0276) and StemFlex media (Gibco A3349401), passaged every four days at 1:20 using ReLesR (STEMCELL Technologies 100-0483) and checked for the undifferentiated morphology. To differentiate hESCs or iPSCs, cells were dissociated as single cells with accutase (STEMCELL Technologies 07920), counted and plated at a density of 10^5 cells/mL on matrigel-coated plates (Corning 354277) with 10 μM Rock Inhibitor Y-27632 (STEMCELL Technologies 72308). All differentiation was performed in a basal media that includes Advanced DMEM/F12 (Gibco 12634010), N2 Supplement (Gibco 17502048), B27 Supplement (Gibco 17504044), and Penicillin-Streptomycin-Glutamine (Gibco 10378016), with the addition of signal agonists and antagonists specified in the following sections.

#### Sequential combinatorial signal screen with pooled scRNAseq

For the hM_d4 combinatorial signal screen, hESCs were plated at 5104 cells/well as single-cells into 24-well cell-culture plates. Plates were coated with matrigel (Corning 354277) at room temperature for an hour before use. After 24 hrs of recovery in StemFlex feeder free media, cells were differentiated into the middle primitive streak (mPS) state with the mPS induction media for 1 day (Step 1). The mPS induction media was supplemented with 6 μM CHIR99021 (Fisher Scientific 44-231-0), 40 ng/mL BMP4 (R&D Systems 314-BP-050/CF), 30 ng/mL Activin A (R&D Systems 338-AC-010/CF), 20 ng/mL FGF2 (R&D Systems 3718-FB-010), and 100 nM PIK90 (EMD Millipore 528117).

The mPS cells were then used for the sequential combinatorial signal screen that differentiated the cells for 3 more days (Step 2 for 1 day + Step 3 for 2 days). Among the combinations, three different lateral plate mesoderm (LPM) states (Day 2) were induced following published protocols ^17,29^. Specifically, mPS was induced into cardiac LPM with 30 ng/mL BMP4 (R&D Systems 314-BP-050/CF), 1 μM A83-01 (Fisher Scientific 29-391-0), and 1 μM WNT-C59 (Fisher Scientific 51-481-0). mPS was induced into anterior foregut (aFG)-LPM with 30 ng/mL BMP4, 1 μM A83-01, 1 μM WNT-C59, 2 μM RA (Fisher Scientific AC207341000), and 1 μM PMA (Fisher Scientific 50-197-5893). mPS was induced into posterior foregut (pFG)-LPM with 30 ng/mL BMP4, 1 μM A83-01, 1 μM WNT-C59, and 2 μM RA. LPM cells were further differentiated into their respective splanchnic mesoderm (SM) fate by cardiac SM induction media (30 ng/mL BMP4, 1 μM A83-01, 1 μM WNT-C59, 20 ng/mL FGF2), aFG-SM induction media (30 ng/mL BMP4, 1 μM A83-01, 1 μM WNT-C59, 20 ng/mL FGF2, 2 μM RA, 1 μM PMA), or pFG-SM induction media (30 ng/mL BMP4, 1 μM A83-01, 1 μM WNT-C59, 20 ng/mL FGF2, 2 μM RA) for 2 days. The rest of the sequential combinatorial conditions were chosen either by random combinations of the five signaling pathways (54 combinations) or bit-flip combinations of the five signaling pathways based on the three controls (38 combinations).

Media was changed every day. On day 4, cells were incubated in Rocki for 30 minutes, dissociated as single cells in accutase, and barcoded with multiSeq oligos (Millipore Sigma LMO-001). Each well in the screen received a unique barcode. Multiplexing was performed in accordance with the manufacturer’s protocol ^28^. Cells were resuspended in PBS + 1% BSA (Sigma Aldrich A9418), pooled and sorted for live cells using a DAPI stain. Cells were processed using a 10X Chromium flow-cell and sequenced on a Illumina Novaseq sequencer, and 19,155 cells passed quality controls.

For the hM_d7 combinatorial screen, cells were plated as and differentiated to the mPS state as in hM_d4. Next, they were differentiated into one of four states for 1 day using the LPM condition (30 ng/mL BMP4, 1 μM A83-01, 1 μM WNT-C59) with or without 2 μM RA and 1 μM PMA. For all future steps, the RA and PMA levels were kept constant. Next, they were differentiated to the splanchnic mesoderm-like state for 2 days with (30 ng/mL BMP4, 1 μM A83-01, 1 μM WNT-C59, 20 ng/mL FGF2). Finally, all possible combinations of 30 ng/mL BMP4, 1 μM A83-01, 1 μM WNT-C59, 20 ng/mL FGF2 were then provided to each of the four states for 3 days, totaling 64 conditions. Cells were pooled and sequenced as described in hM_d4. Cells were processed as in hM_d4, and 11,857 cells passed quality controls.

For the hE_d8 combinatorial screen, cells were plated using the same procedure described in hM_d4 with a density of 2 x 10^-5^ cells/mL. They were first differentiated to the anterior primitive streak (APS) for 1 day (100 ng/mL Activin A, 2 μM CHIR), which was then differentiated into the definitive endoderm (DE) for 2 days (100 ng/mL Activin A). After this, all combinations of 30 ng/mL BMP4, 1 μM A83-01, 1 μM WNT-C59, 20 ng/mL FGF2, 2 μM RA and 1 μM PMA were then provided for 5 days to create 64 combinations. Cells were pooled and sequenced as described in hM_d4. Cells were processed as in hM_d4, and 9,921 cells passed quality controls.

Count matrices were generated using CellRanger (v7.1.0) and aligned to the reference human genome GRCh38. Barcodes were aligned and demultiplexed using the R library deMULTIplex (version 1.0.2) ^28^. Barcodes were visually inspected using tSNEs of cells clustered in barcode space. This was performed in accordance with the criteria specified in deMULTIplex. Once a list of observed barcodes was identified, each barcode was normalized to the number of barcodes observed per cell. Each barcode then had outliers trimmed and its probability density function computed. Lastly, thresholds were identified by defining the local maxima, with the mode for negatives and maximum for positive values. Barcodes were then assigned independently for each cell. Cells with one barcode are designated as singlets, more than one as doublets, and those with none as negatives. The optimal threshold was chosen which maximized the number of singlets relative to doublets and negatives. 65, 55 and 55 unique barcodes were recovered in the hM_d4, hM_d7 and hE_d8 screens respectively. Across each of these screens the maximum number of cells per barcode was 975, 1255, 807, respectively.

#### Directed differentiation into respiratory mesenchyme

hESCs or iPSCs were first differentiated into aFG-SM using the aforementioned media condition. Then they were induced with either IRIS-optimized media protocol (30 ng/mL BMP4, 2 μM RA, 2 μM PMA, 3 μM CHIR) for 3 days, or published protocol (30 ng/mL BMP4, 2 μM RA, 2 μM PMA) for 3 days with 2 μM CHIR added only for the last day ^29^. On Day 7 of differentiation, samples were fixed in 4% PFA for 15 minutes, then processed for HCR.

#### Mouse foregut explant culture

To perform the signaling perturbation in explants, E9.0 mouse embryos were harvested from wild type C57BL/6J timed matings. Foreguts were isolated by dissecting the anatomical region between the pharyngeal arches and liver, excluding additional tissues including the somites. Foreguts were cultured in DMEM/F12 (Gibco 10565042) + 5% FBS (Takara 631368) + 1% Penicillin/Streptomycin/Glutamate (Gibco 10378016) supplemented with either 3 μM of CHIR or 1 μM C59. After 24 hours in culture at 37℃, samples were fixed in 1% PFA overnight at 4℃ and processed for subsequent analysis.

#### Hybridization Chain Reaction (HCR)

HCR-FISH probes were designed in-house and tested on samples with known expression to validate probe efficiency ^54^. Mouse foregut explant or differentiated hESC samples were dehydrated with 100% methanol and subsequently processed according to the HCR-FISH protocols from Molecular Instruments. In brief, samples were hybridized overnight at 37℃ with 4 μM of probes and amplified overnight at room temperature with B1, B3, B4, B5 hairpin amplifiers (Molecular Instruments). Afterwards, hESC-derived samples were imaged on a Nikon TI2 widefield fluorescence microscope. Explant samples were imaged on a Zeiss LSM980 Airyscan 2 confocal microscope. Analysis of the hESC-derived cells was performed by first segmenting in cellpose using the ‘cyto2’ model (parameters: diameter = 35, flow-threshold = 0.7) ^55^. Total nuclear fluorescence signal was then measured in each channel with these segmentation masks. Individual fluorescence measurements for each cell were grouped based on the treatment condition and a student’s T-test was performed to determine statistical significance using scipy.stats.ttest_ind (version 1.12.0). This test assumes the expression level following a T-distribution and values are considered significant if p < 0.05. Data was visualized using seaborn.violinplot.

## Code Availability

Code is available at https://github.com/Pulin-Li-Lab/IRIS-signaling-inference.

## Data availability

All data analyzed in this article is available publicly through online sources. The following data are available at the Gene Expression Omnibus database with their corresponding accession numbers: raw data for hM_d7, hE_d8 and hM_d4 (GSE289836), the mESC screen (GSE122009), mouse foregut organogenesis dataset (GSE136689), and human airway epithelium (GSE246368). The gastrulation atlas is available on ArrayExpress (E-MTAB-6967).

**Extended Data Figure 1.**
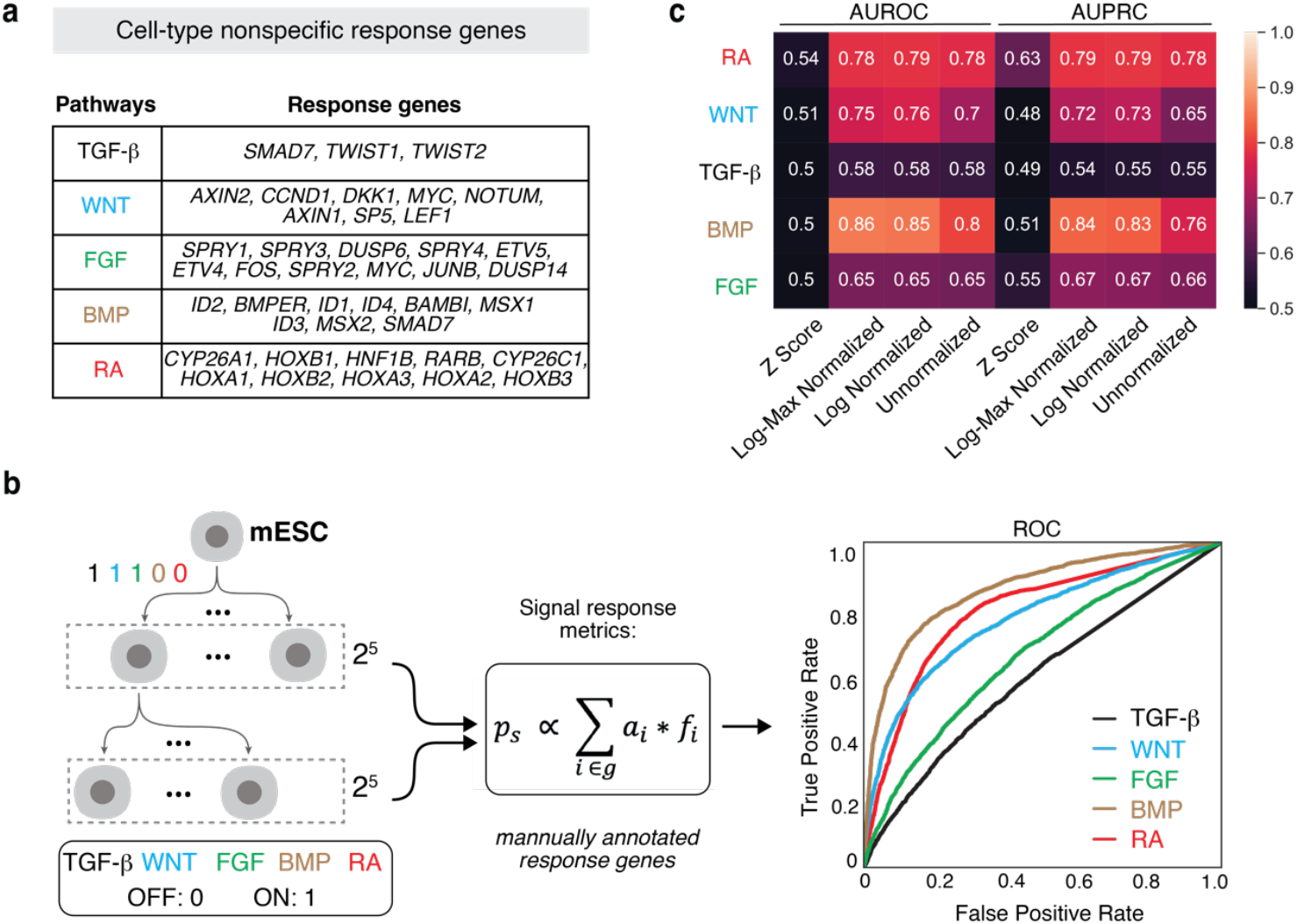
Signal inference using the response-gene method. **a**, A list of response genes chosen for each signaling pathway was modified from Han *et al*. 2020 ^17^. **b**, Response genes inferred signaling pathway activities with limited accuracy, as tested with a ground-truth dataset of mESC differentiation ^21^. Signaling states are represented as binary strings. The probability of being a responding cell (*p*_*s*_) was computed as summed expression of response genes (*f*_*i*_) with weights (*a*_*i*_). ROC, receiver operating characteristic. **c**, The performance of the response-gene method for signal inference, computed based on different normalization strategies for gene expression. AUROC, area under the receiver operating curve; AUPRC, area under the precision recall curve.

**Extended Data Figure 2.**
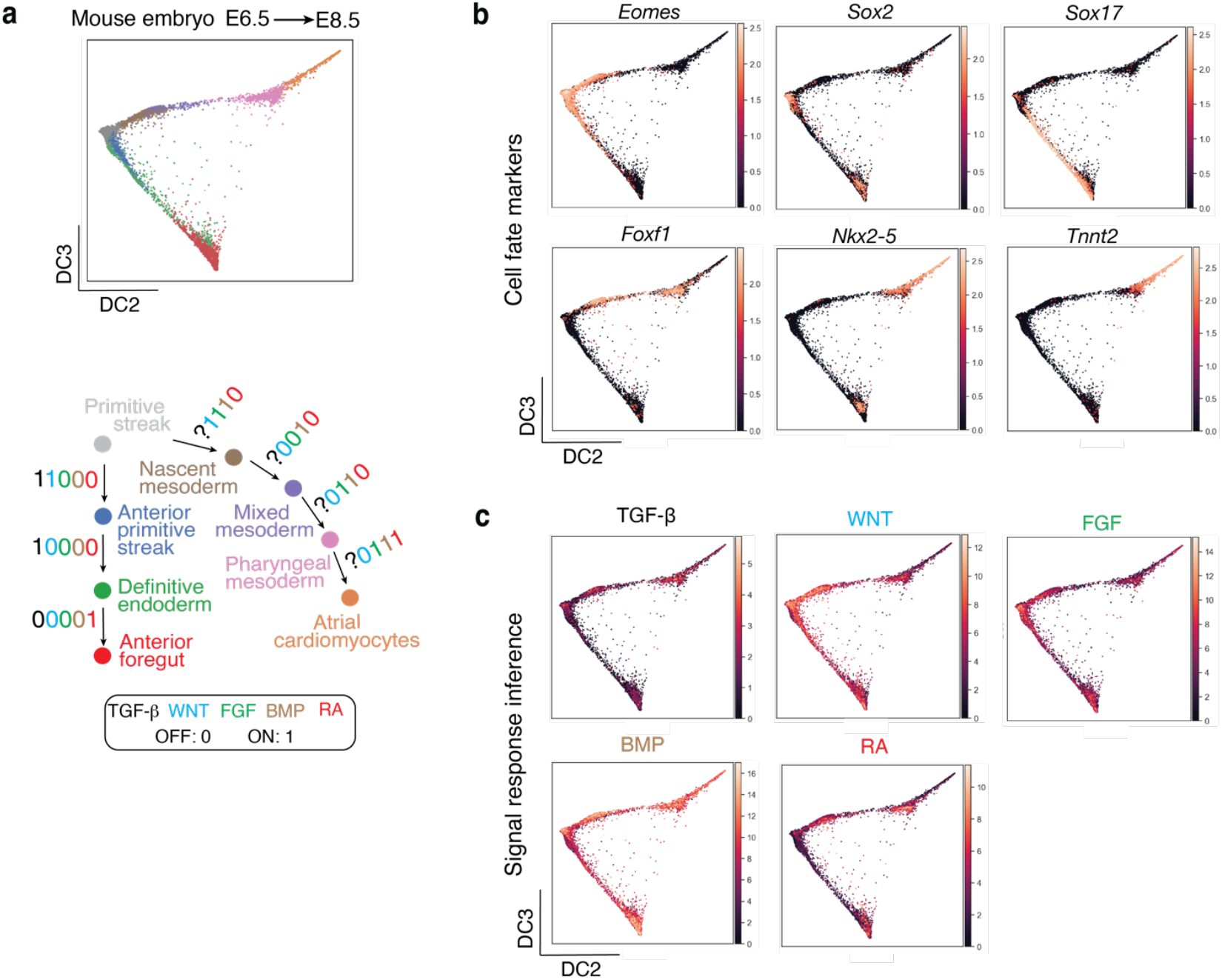
Signal inference of *in vivo* cell types using the response-gene method. **a**, Construction of the anterior foregut and atrial cardiomyocyte trajectories, using cell populations from the mouse embryo (E6.5-E8.5) scRNAseq dataset ^35^. The colors of the cell populations in the diffusion map correspond to those in the schematic. Signal combinations at each step of differentiation were annotated based on the literature. Due to lack of knowledge in the literature, the activity of TGF-β along the cardiac lineage was left out in the annotation. **b**, The expression of key cell fate markers associated with each of the two lineages (normalized to counts per 10^4^). **c**, Signal response-gene scores for all five signaling pathways. Consistent thresholds for binary classification from these scores are difficult to calibrate, limiting their interpretability.

**Extended Data Figure 3.**
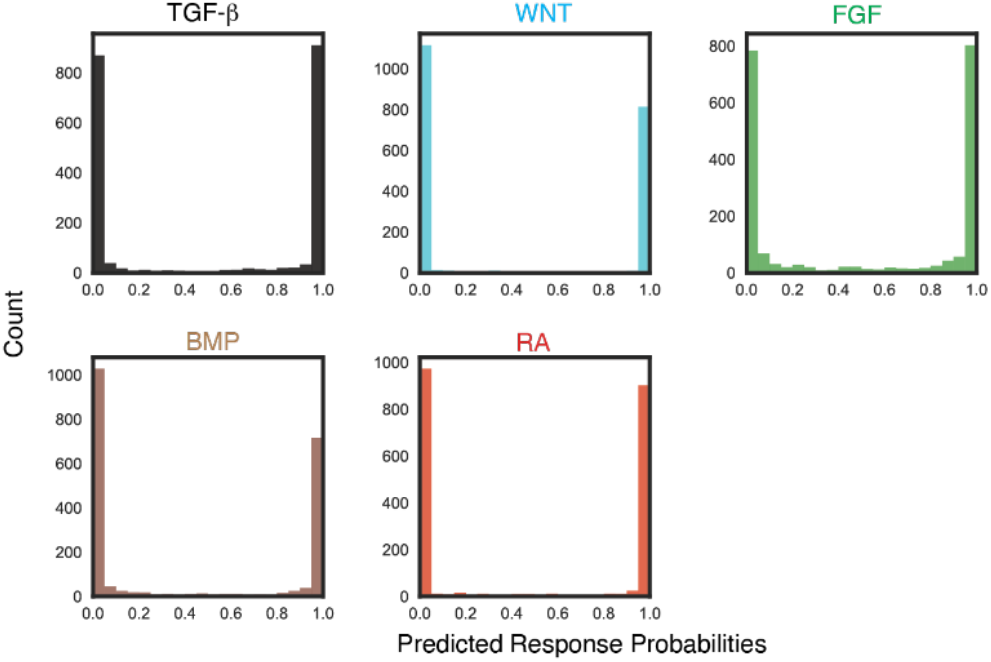
Distribution of IRIS predicted response probability and thresholding. IRIS predicted response probability values were close to 0 and 1 in the in-sample test using mouse screen data, potentially due to the saturation concentrations of ligands used in the screen. 0.5 was chosen as the threshold for the model to be agnostic to the types of downstream errors.

**Extended Data Figure 4.**
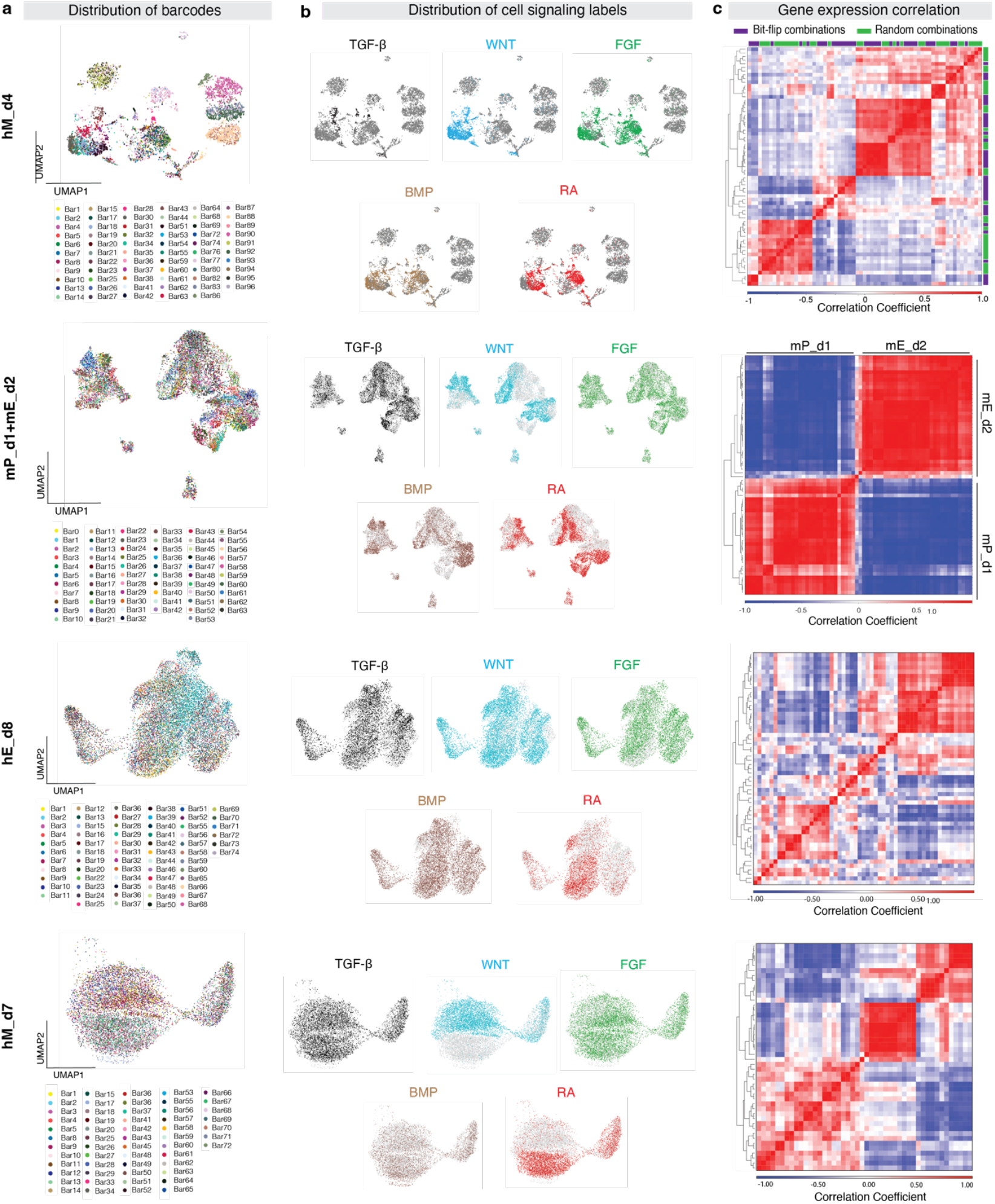
Analysis of cell-state diversity in the mESC and hESC screens. **a**, Distribution of barcodes recovered after sequencing for each screen. **b**, Cells activated with the respective signaling pathways in the last step of differentiation are highlighted by the corresponding color. Step 2 or Step 3 (hM_d4), Step 4 (hM_d7), and Step 3 (hE_d8). **c**, The correlation coefficient averaged across the transcriptome between distinct signal combinations within each screen, highlighting the diversity of final cell states.

**Extended Data Figure 5.**
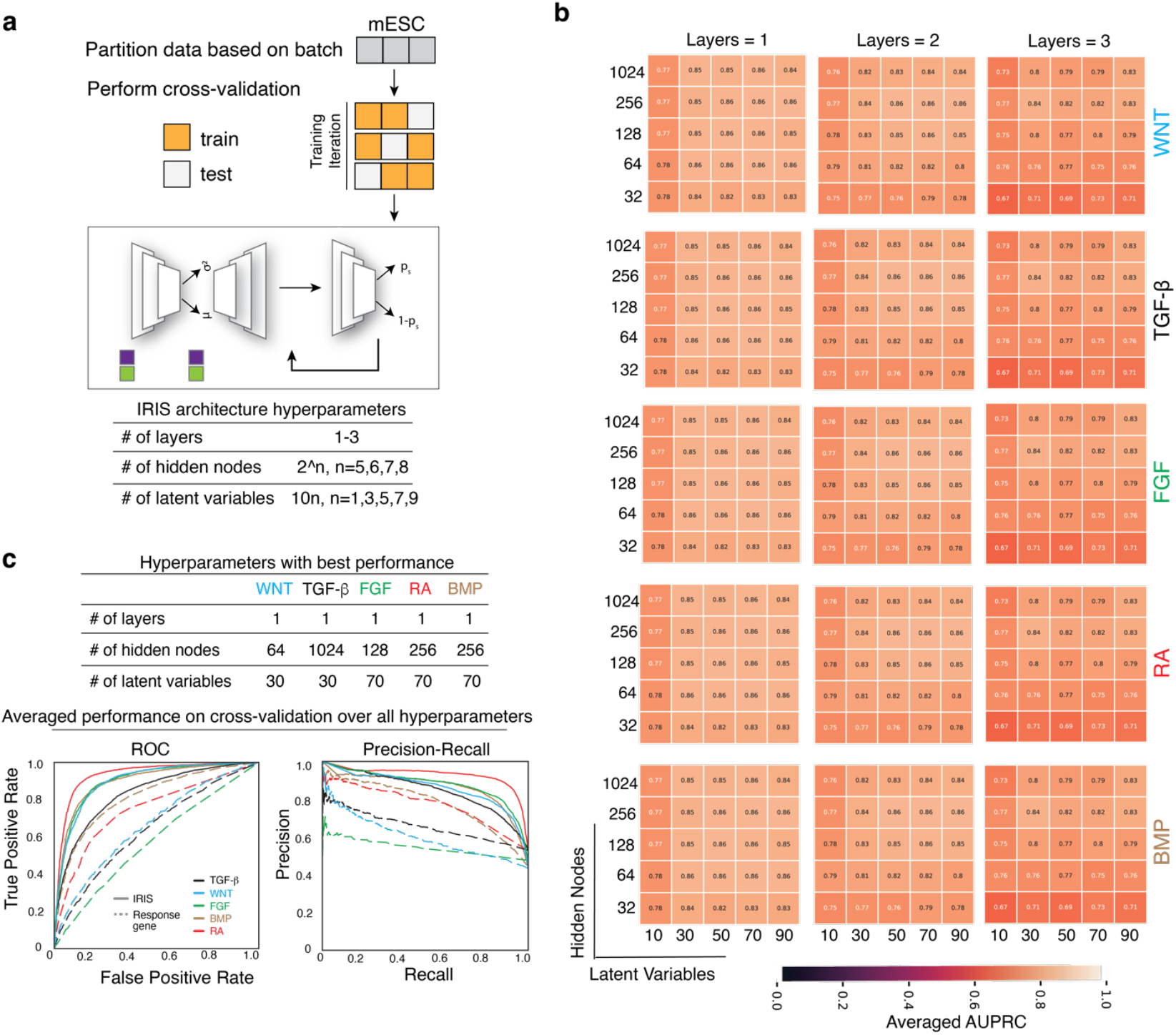
A hyperparameter screen of IRIS reveals optimal architecture for each pathway. **a**, Schematic of hyperparameter optimization. The mESC screen was partitioned into three batches during data collection. In the hyperparameter screen that aims to optimize for cross-batch generalizability, two of the three batches were used for training at a time and the remaining batch was used as a test. Multiple parameters were screened for layers, hidden nodes, and latent variables. **b**, The performance for each combination of the tested hyperparameters was evaluated based on the AUPRC (Area Under the Precision-Recall Curve) averaged across all three training iterations. **c**, Averaged cross-batch performance metrics of the ideal hyperparameters. The optimal parameters were chosen from the hyperparameter screen and all pathways showed improvement over the response gene approach. Performance was averaged across all three iterations.

**Extended Data Figure 6.**
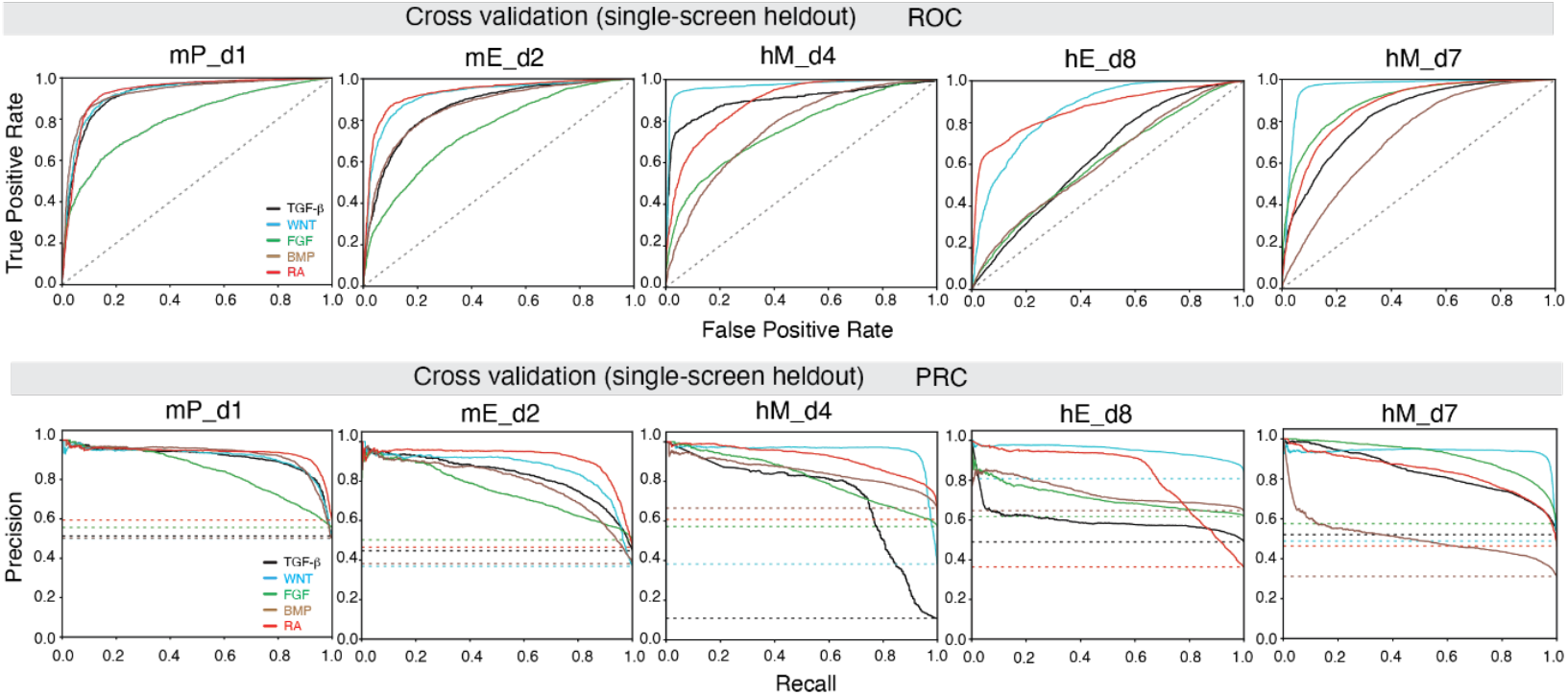
Model evaluation via a cross-validation test. The AUROC (*top*) and PRC (*bottom*) curves of the cross-validation test, with a single screen held out each time (related to Fig. 2c). A random baseline for each pathway (*dashed lines*) was used for comparison.

**Extended Data Figure 7.**
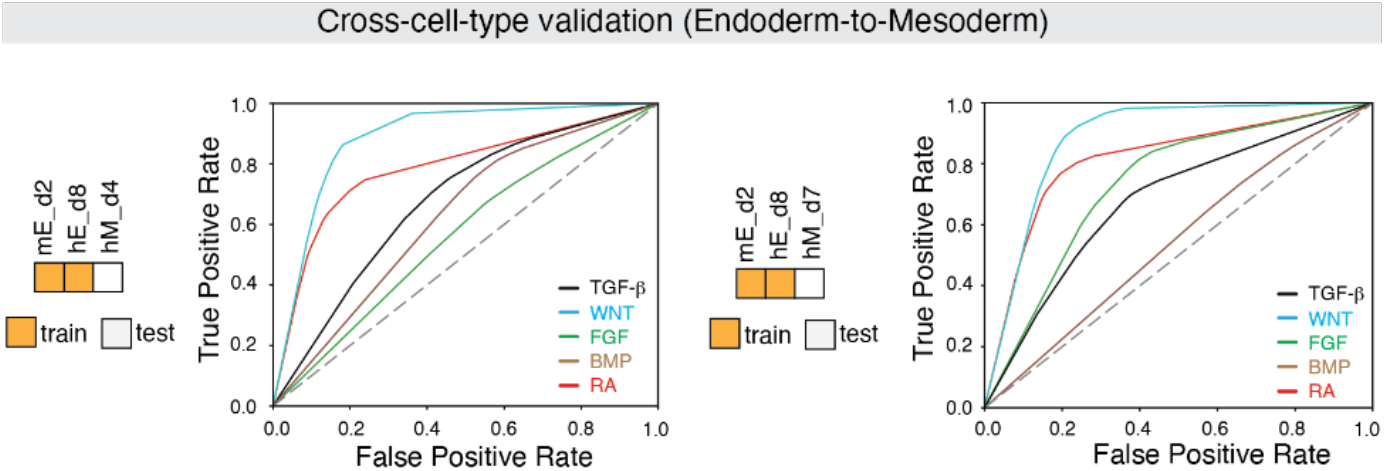
Evaluating cross-cell-type generalization. The AUROC curves of the cross-cell type generalization test, in which IRIS was trained on two endodermal screens (mE_d2, hE_d8) and tested on mesodermal screens (hM_d4, hM_d7) (related to Fig. 2d)

**Extended Data Fig 8.**
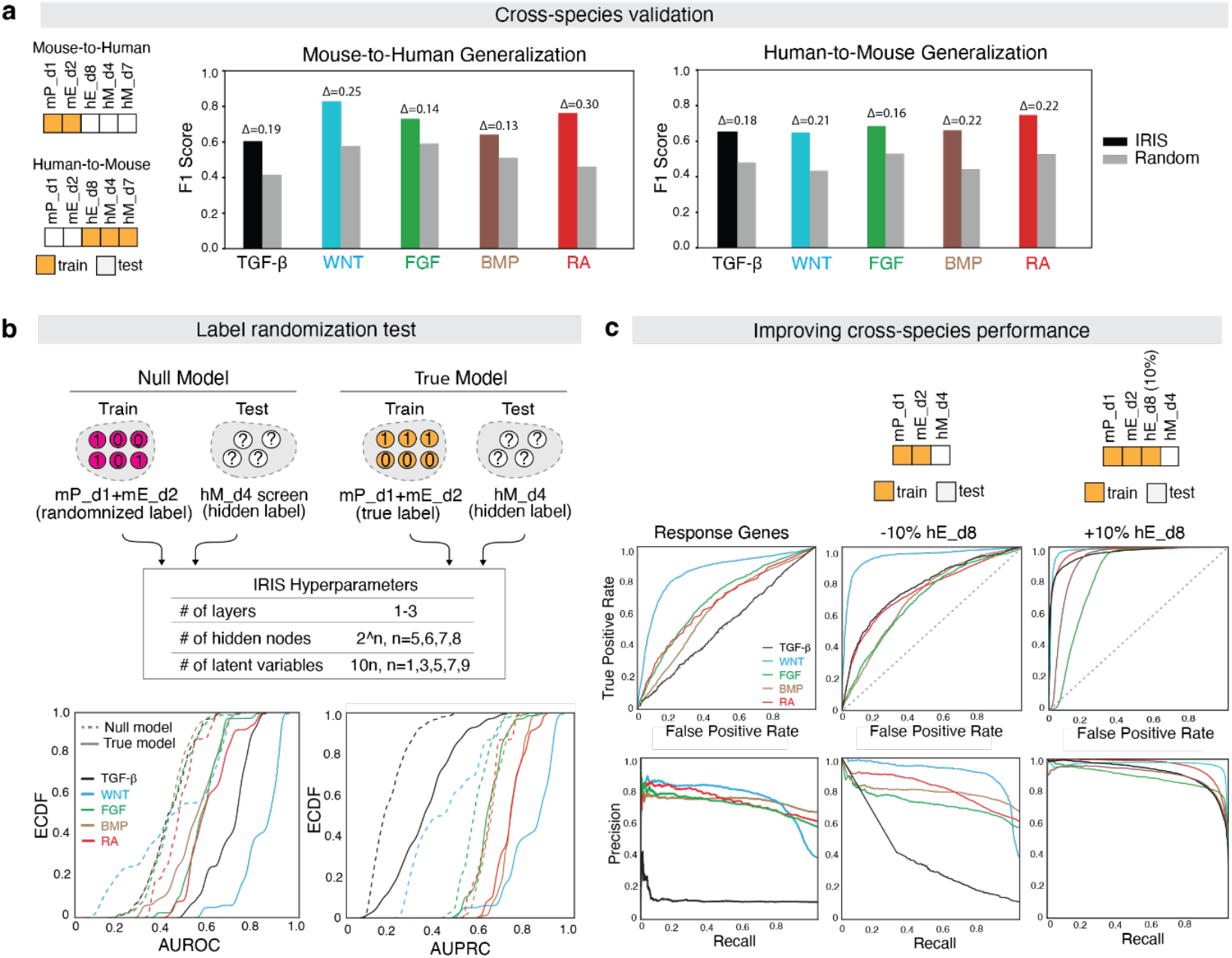
Cross-species generalization test. **a**, Mouse-to-human and human-to-mouse generalization tests, in which models were trained exclusively on mouse screens (mP_d1, mE_d2) and evaluated against all human screens (hM_d4, hM_d7, hE_d8), or vice versa. The performance was compared to random baselines. **b**, Cross-species label randomization test. Comparison between a Null Model and a True Model, both trained exclusively on mouse screens and evaluated for their inference accuracy on the hESC screen (hM_d4) across a range of hyperparameters. The True Model (*solid lines*) consistently outperforms the Null model (*dashed lines*), suggesting IRIS is learning meaningful response features in the cross-species context. ECDF, empirical cumulative distribution function. AUROC, area under the receiving operator curve. AUPRC, area under the precision recall curve. **c**, Evaluating the effect of including in-species (but cross-cell-type) training data on cross-species performance. IRIS was trained either exclusively on the mouse screens or together with 10% of hE_d8, to predict on the unrelated hM_d4 screen.

**Extended Data Figure 9.**
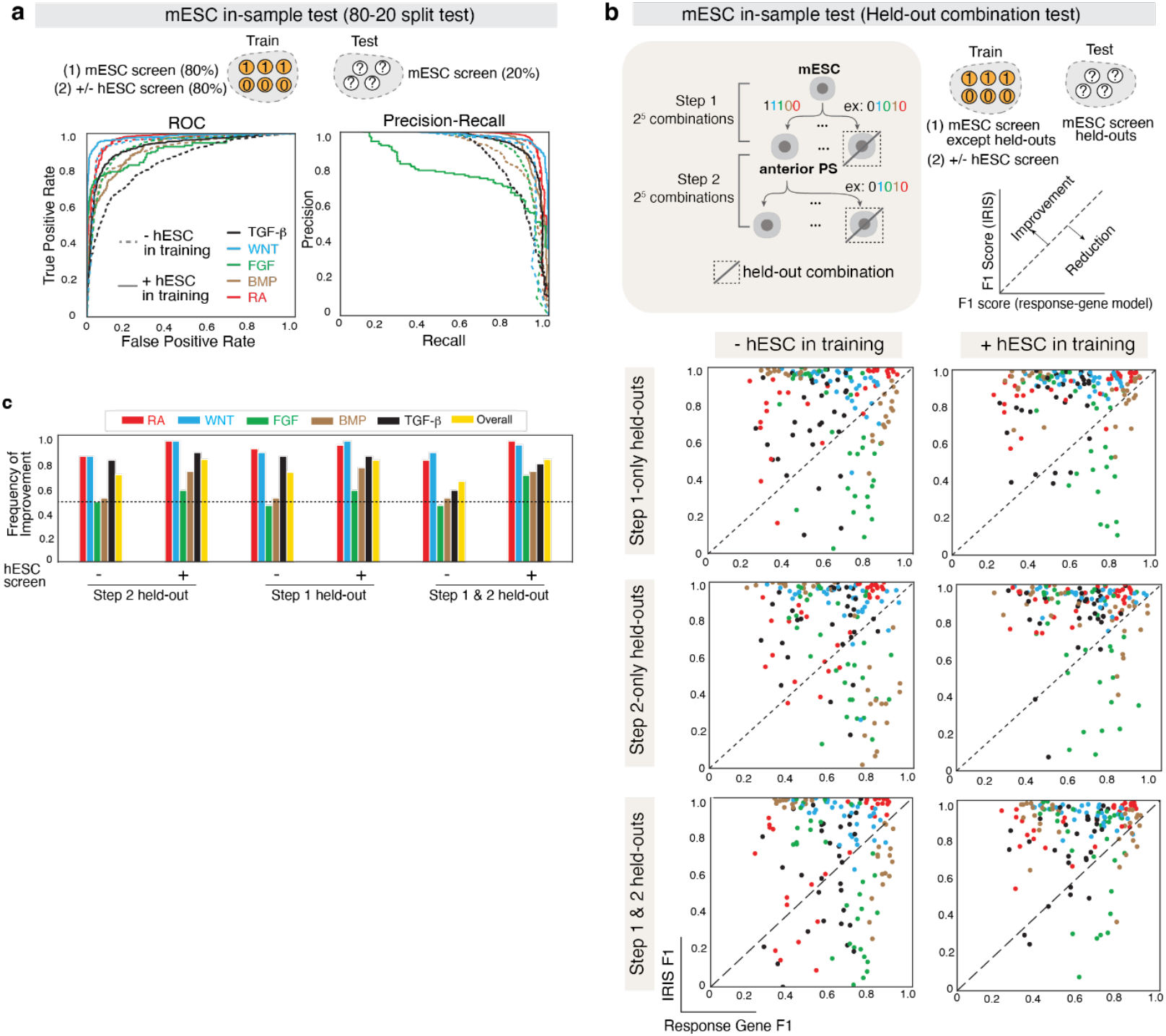
Increasing diversity of cell types in training data broadly improves inference accuracy. **a**, Including the hM_d4 screen in the training data improved in-sample inference accuracy. IRIS was trained on 80% mESC screens with or without 80% of the hM_d4, and tested on the remaining 20% of the mESC screens. **b, c**, Held-out signal combination test showed consistent improvement of IRIS over the response-gene method and including hM_d4 in training further improved the accuracy of IRIS. Samples with the same combination in Step 1 and Step 2 were held out together or separately. Comparison of IRIS and Response Gene F1 scores for each pathway. Individual points represent the performance of IRIS on each signaling pathway in a given held-out condition (**b**). The frequency of improvement over the response-gene method for each signaling pathway was calculated and presented as barplot (**c**).

**Extended Data Figure 10.**
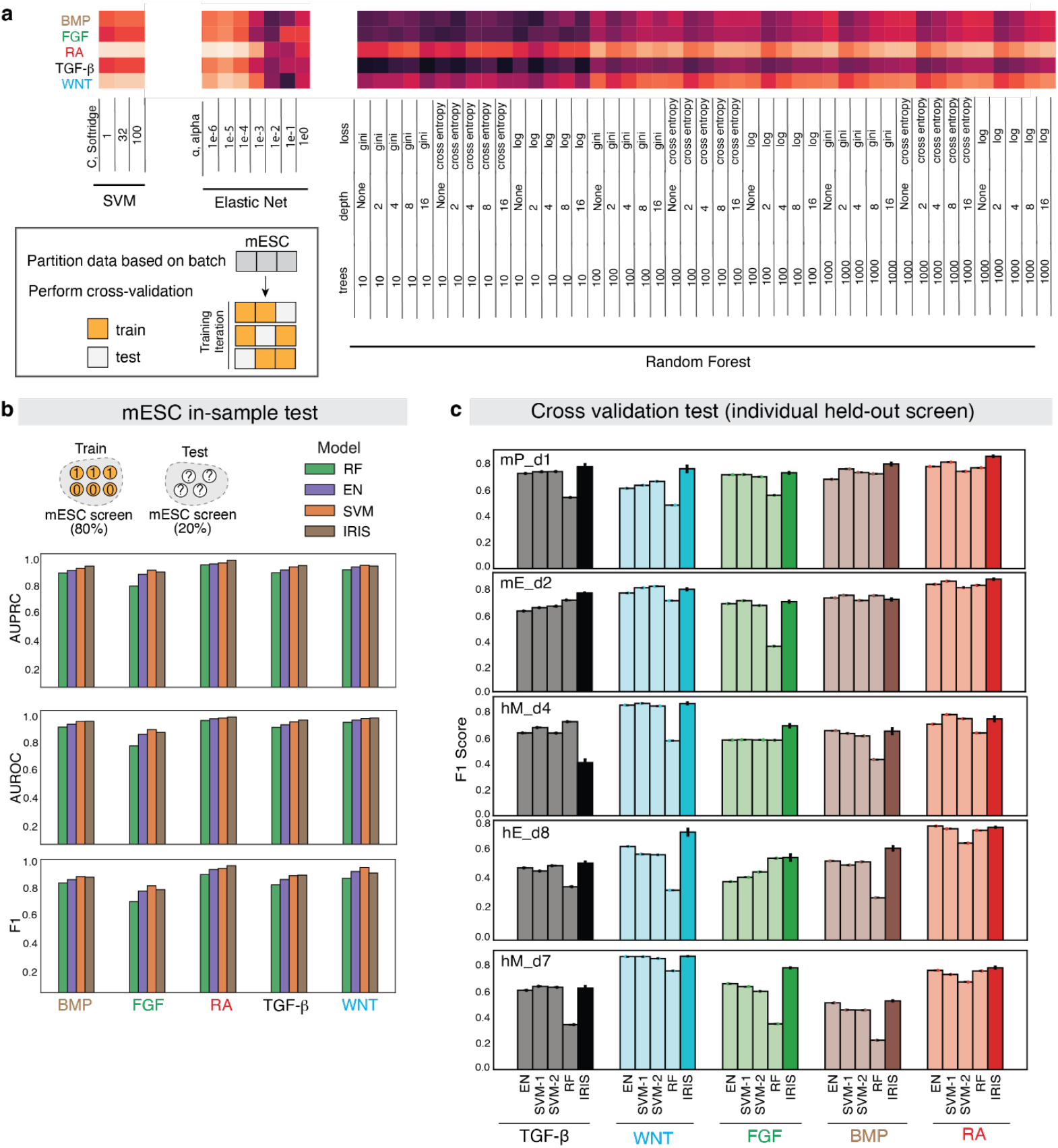
Model benchmarking and hyperparameter optimization for white-box machine learning models. **a**, PCA was first performed on the data in accordance with Scanpy’s default parameters. The mESC screen was partitioned into three batches during data collection. Individual batches were held out one at a time and the average AUPRC is presented in the heat map for a variety of hyperparameters for each model: SVM (softridge, C=1, 32, 100), Elastic net (alpha, 10^-n, n=1, 2, 3, 4, 5, 6), and Random forest (loss=gini, cross entropy, log; number of trees= 10, 100, 1000; max depth=2, 4, 8, 16, None). **b**, Model benchmaking on the in-sample 80-20 split test with the default parameters of each model (related to Fig. 1c). IRIS outperforms random forests (p < 0.05; one-sided binomial test, n=5) but there was no statistically significant difference between IRIS and SVM or EN. **c**, Model benchmarking via a cross-validation test (related to Fig. 2c). IRIS is the most consistently performing model across all signaling pathways and splits (p < 0.05; one-sided binomial test on all possible pairwise comparisons against all other models, n=100).

**Extended Data Figure 11.**
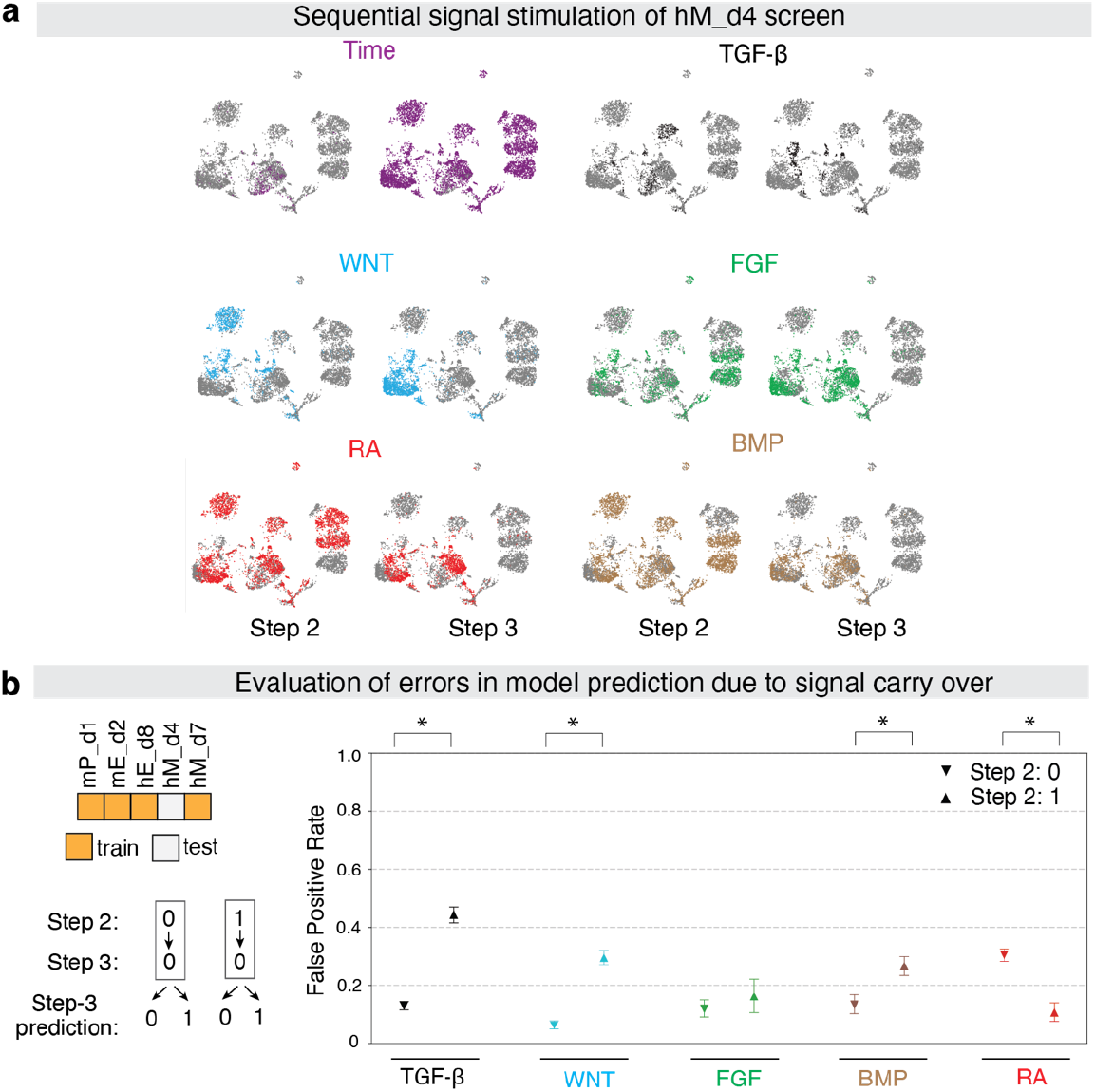
Evaluation of signaling carry-over across time steps. **a**, Ground-truth signaling labels for the hM_d4 screen. Cells with the signaling pathway activated during Step 2 (*left*) or Step 3 (*right*) are shown separately, and they are highlighted in the corresponding color on the UMAP plots. The “Time” UMAP plots highlight the cells that were sequenced after Step 2 or Step 3 stimulation separately. **b**, Evaluation of false positive rates (FPR) in model prediction due to signal carry-over. For each signaling pathway, the FPR in the Step 3 was calculated for cells that had received the same signal in Step 2 (*right*) and compared against cells that had not received the same signal in Step 2 (*left*). Confidence intervals were determined using bootstrapping of n=1000 samples. Significance was determined via a z-test with standard errors (n=3903; * indicates p < 0.05).

**Extended Data Figure 12.**
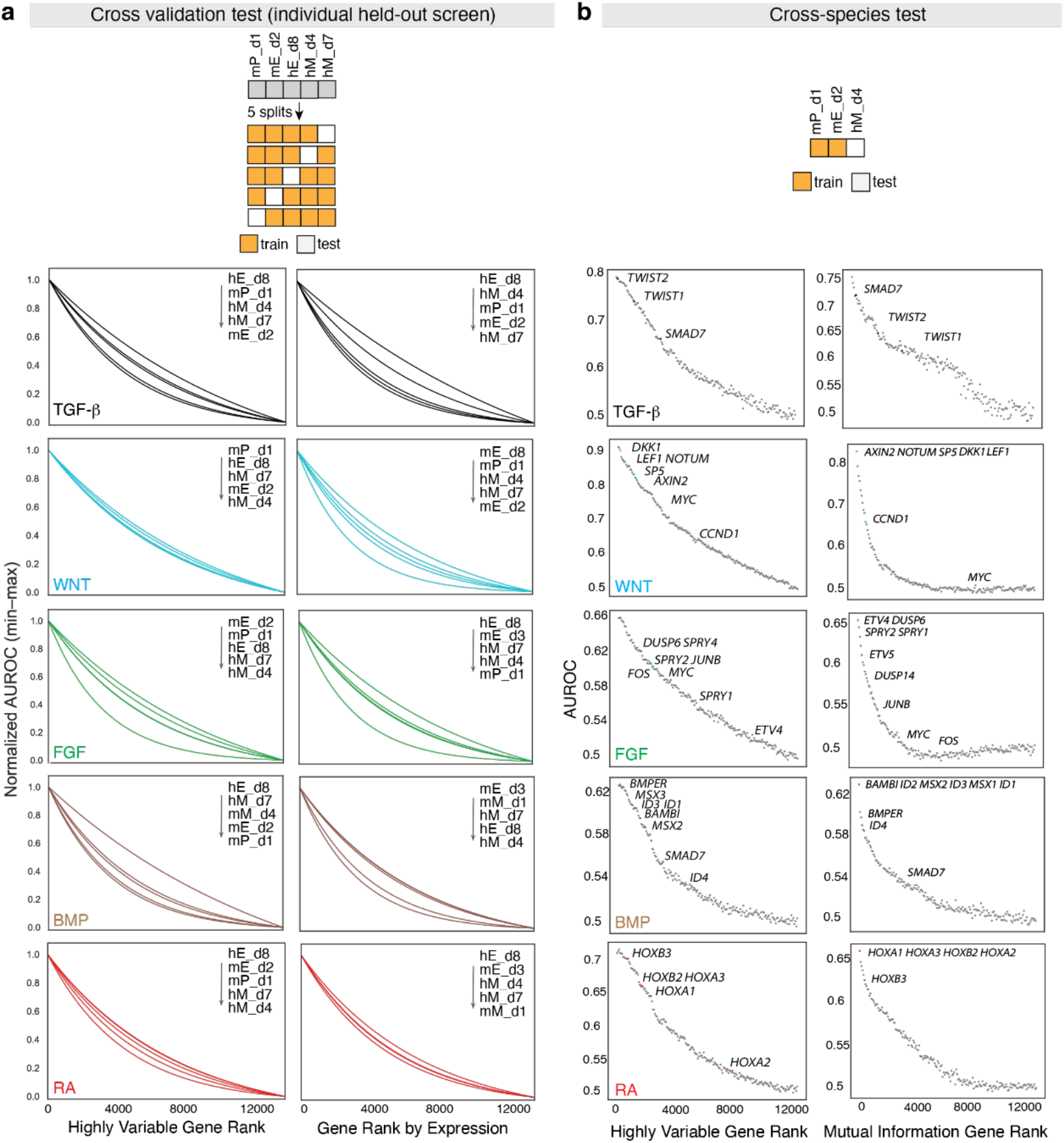
Gene ablation tests. **a**, Gene ablation test on the single screen-heldout cross-validation (related to Fig. 2f). Genes were ranked according to either the highly variable (*left column*) or highly expressed (*right column*) gene metrics. The inference accuracy of IRIS was computed after each round of corruption. The result for each data split and each pathway was fit to an exponential function and normalized between 0 and 1 for visualization. AUROC declines gradually over thousands of ranked genes. **b**, Cross-species gene ablation test. Genes were ranked according to either the highly variable (*left column*) or mutual information (*right column*) gene metrics. Dots represent individual groups of genes for visualization. The points which correspond to manually annotated response genes (Extended Data Fig. 1) are highlighted.

**Extended Data Figure 13.**
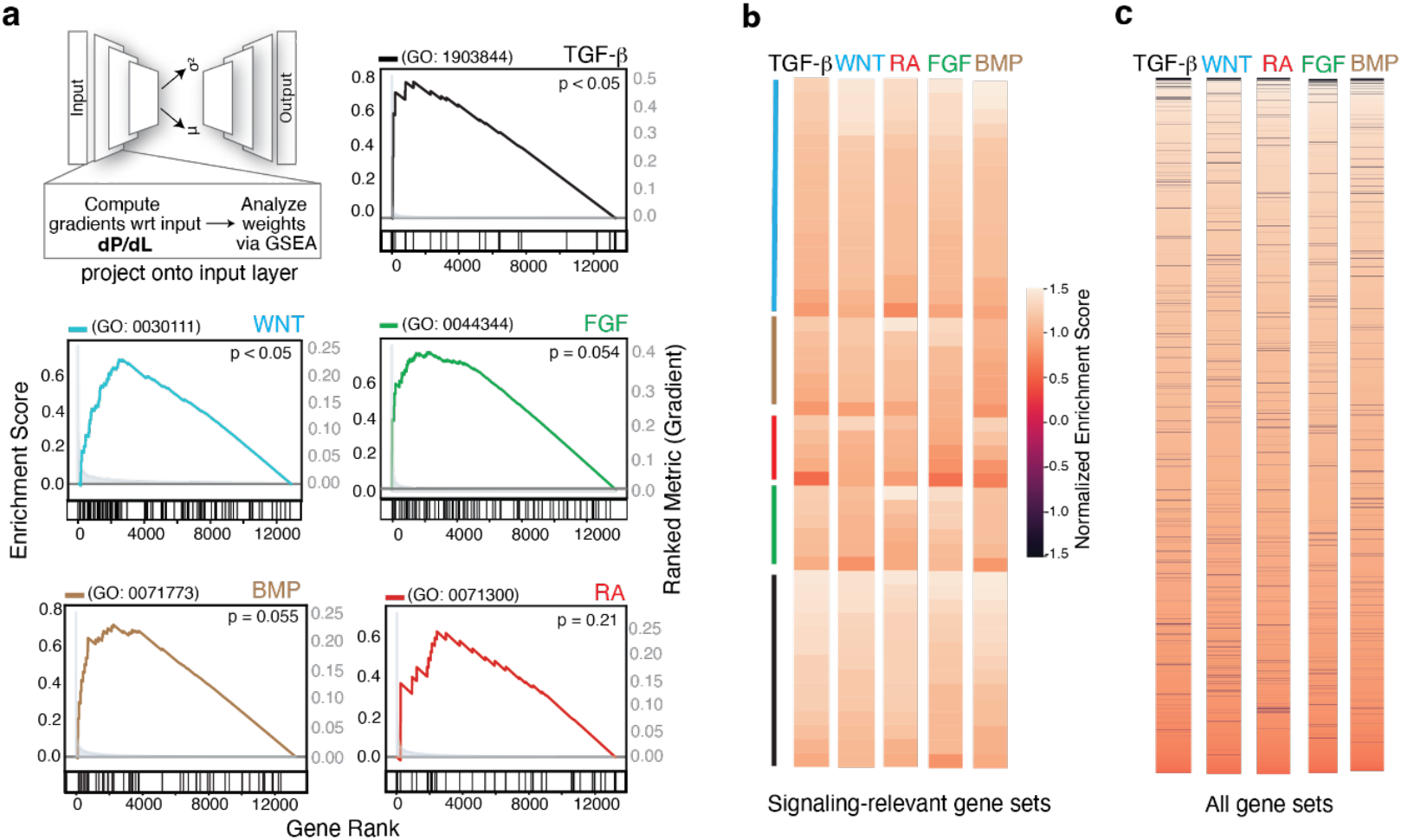
Gene saliency test. **a**, To estimate the contribution of each gene to the model performance, gradients of the loss with respect to the input variables (dP/dL) were computed for each pathway during training. Genes were ranked based on the gradient and a GSEA (Gene Set Enrichment Analysis) was performed on each pathway by choosing the gene set annotated as “response to” the corresponding signal (GO number labeled). The p-values were computed with a permutation test (n=1000). **b**, GSEA on all signaling-relevant gene sets in the database showed no preferential enrichment of the gene sets annotated to be related to the corresponding pathway. Color lines denote the gene sets assigned to the corresponding signaling pathway by the database. **c**, GSEA for all gene sets present in the GO Biological Pathways 2021 database showed broad enrichment of many gene sets for each pathway.

**Extended Data Figure 14.**
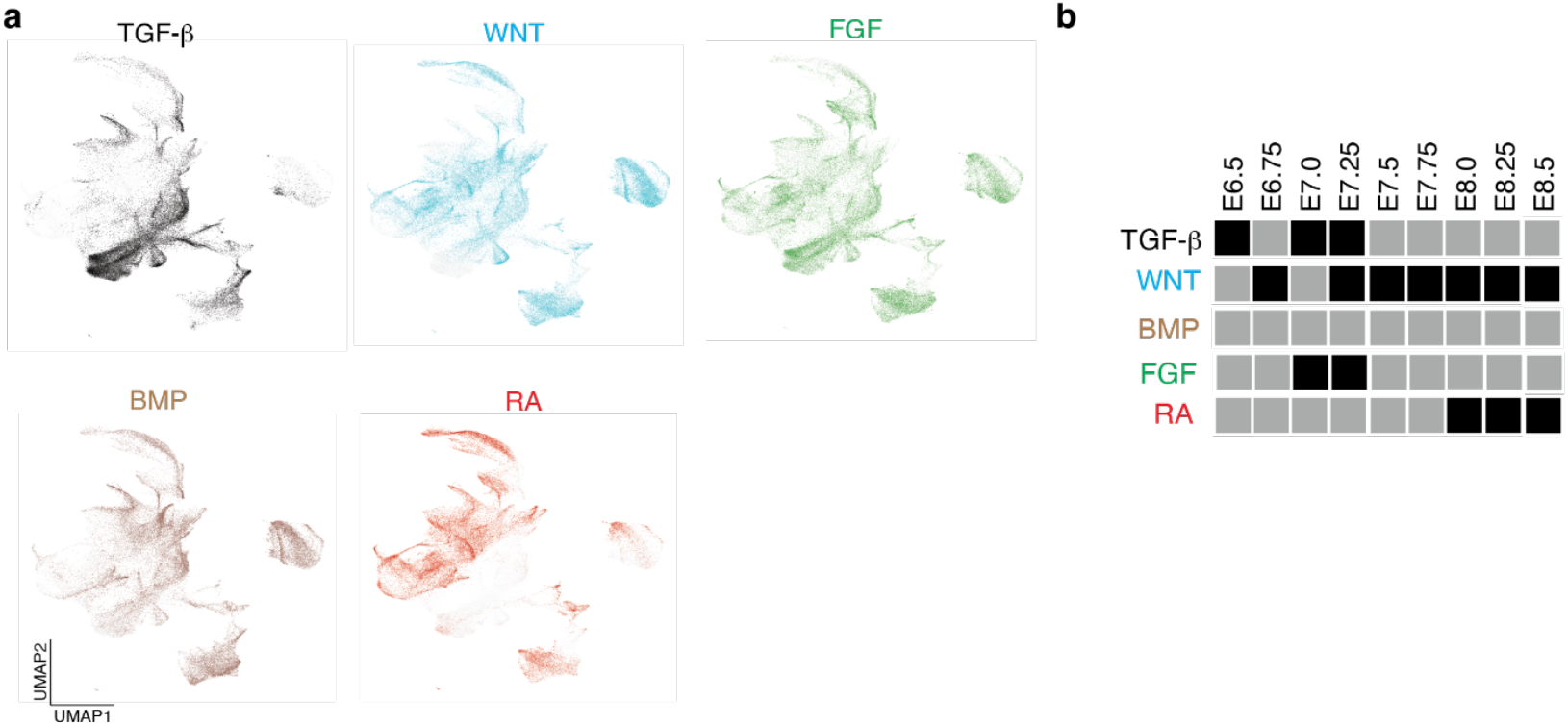
Inferring the activity of individual signaling pathways across the mouse gastrulation dataset. **a**, The five signaling pathway activation predictions annotated on the mouse gastrulation atlas revealed patterns of response across diverse cell types. Cells inferred to have the denoted signaling pathway activated are highlighted in the corresponding color. **b**, Enrichment of signaling pathway activation across stages of embryonic development. Black squares denote significant enrichment via a one-tailed Mann Whitney u-test (n=139331; p < 0.05) (alternative: greater).

**Extended Data Figure 15.**
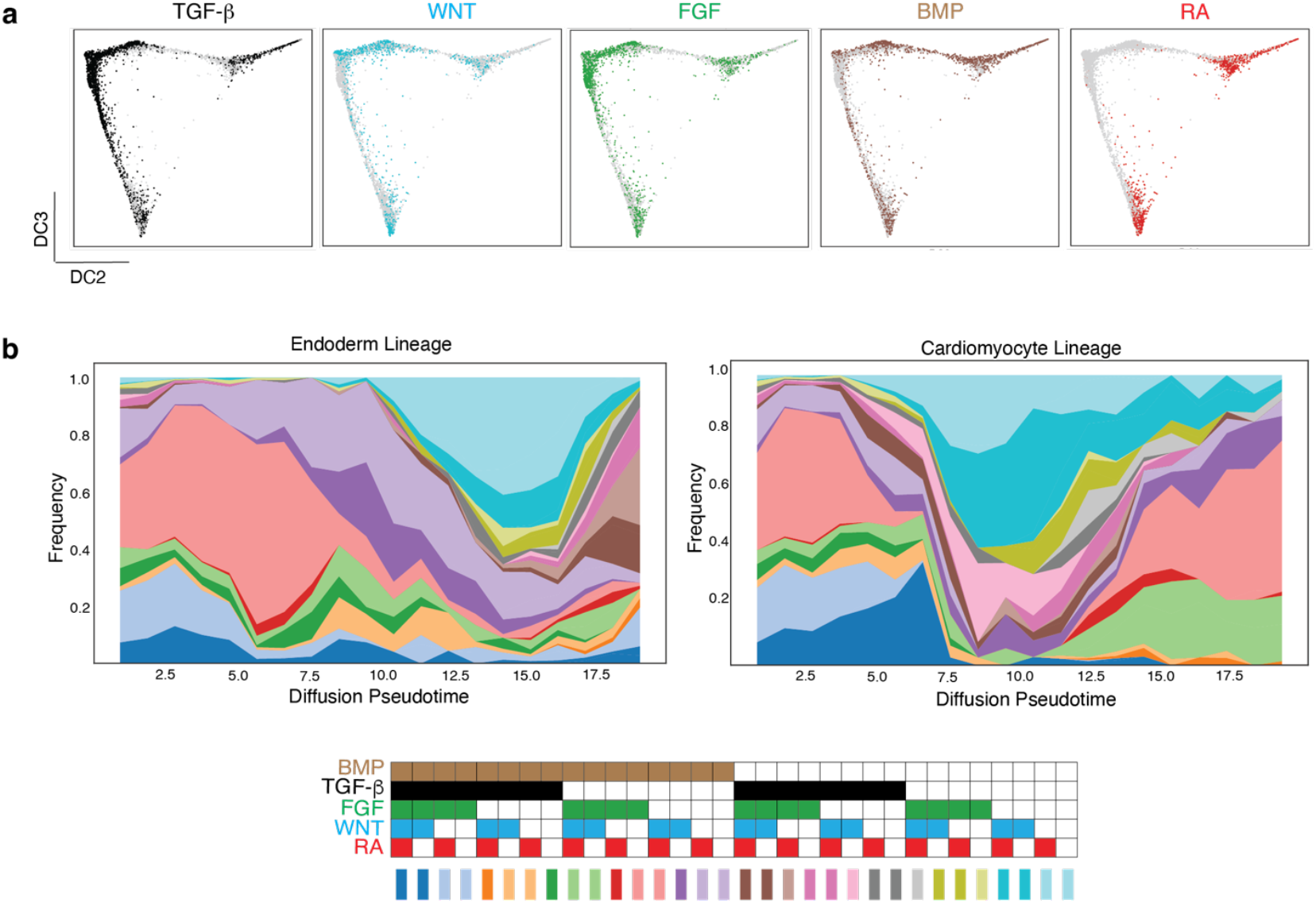
IRIS prediction for the endodermal and cardiac lineages. **a**, Inferred signaling states for individual pathways across both the endodermal and cardiac lineages. Cells are arranged in the same way as they are in **Fig. 3c**. Cells inferred to have the denoted signaling pathway activated are highlighted in the corresponding color. **b**, Frequencies of all inferred signal combinations among the 5 pathways across the inferred pseudotime.

**Extended Data Figure 16.**
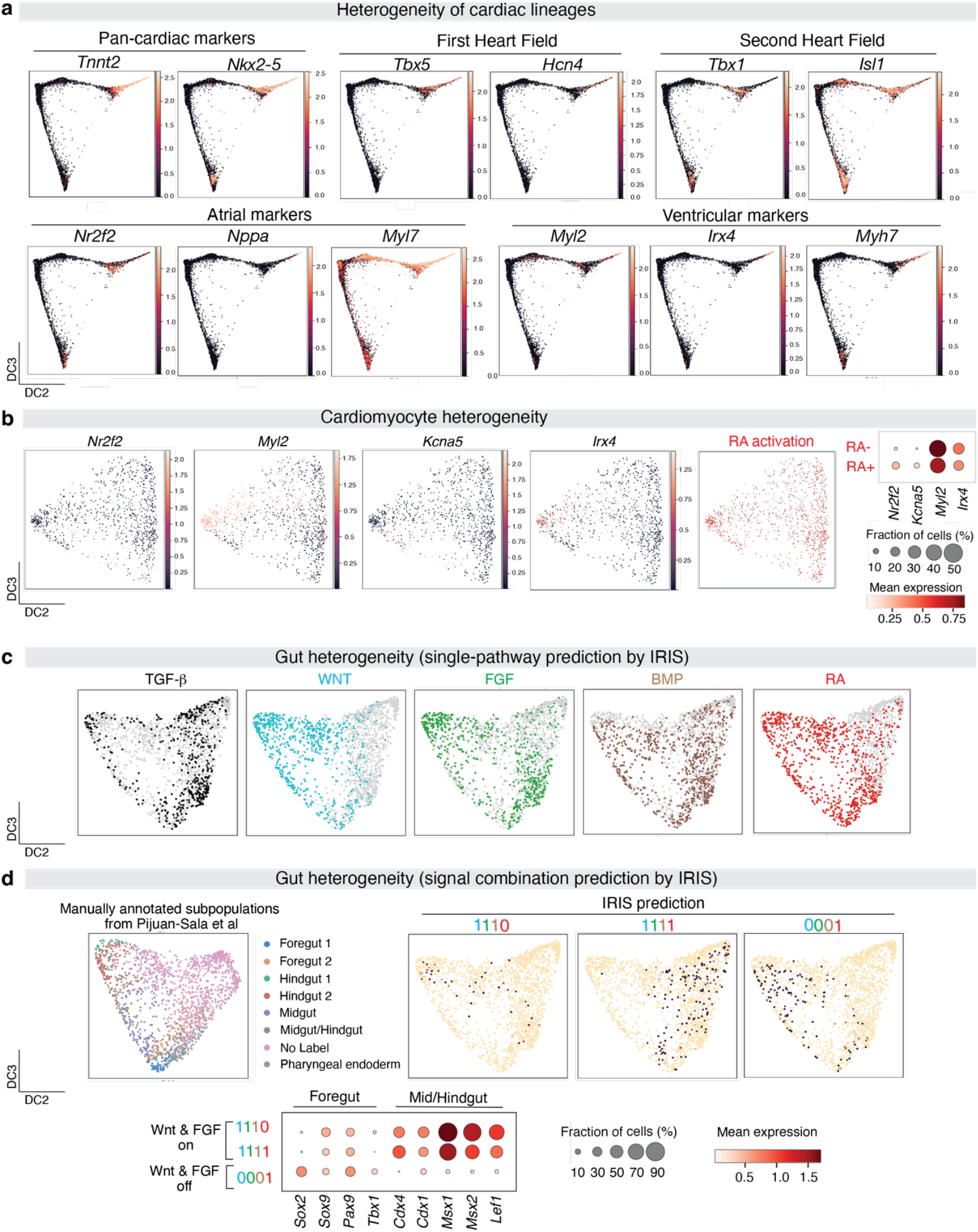
Inferred signaling state heterogeneity correlates with biologically meaningful cell-type heterogeneity. **a**, Markers associated with different developmental stages of cardiac mesoderm align well with the diffusion component position. Both atrial and ventricular markers are expressed in the annotated cardiomyocyte population, revealing the heterogeneity of cell types. **b**, IRIS revealed heterogeneous states of RA activation within the annotated cardiomyocyte cluster (n=1206). Cells predicted to be RA+ were enriched for atrial markers (*Nr2f2, Kcna5;* p < 0.05, one-tailed Mann-Whitney U-test (alternative: greater)), compared to cells predicted to be RA-. **c**, IRIS predicted activation of individual signaling pathways in the annotated gut population. **d**, IRIS identified several main combinatorial signaling states that corresponded to cell fate states within the gut cluster. The gut population (n=1940) was re-clustered and annotated by the original paper based on cell fate markers (*upper left panel*) ^35^. Cells inferred to have the denoted combinations are highlighted (*upper right panel, black dots*). Gut cells associated with different combinatorial signaling states are enriched for different cell-fate markers that correspond to the foregut or mid/hindgut (*bottom*) (*Foregut: Sox2, Tbx1*; *Mid/Hindgut*: *Cdx4, Cdx1, Msx1, Msx2, Lef1*; p < 0.05, one-tailed Mann-Whitney U-test (alternative: greater)).

**Extended Data Figure 17.**
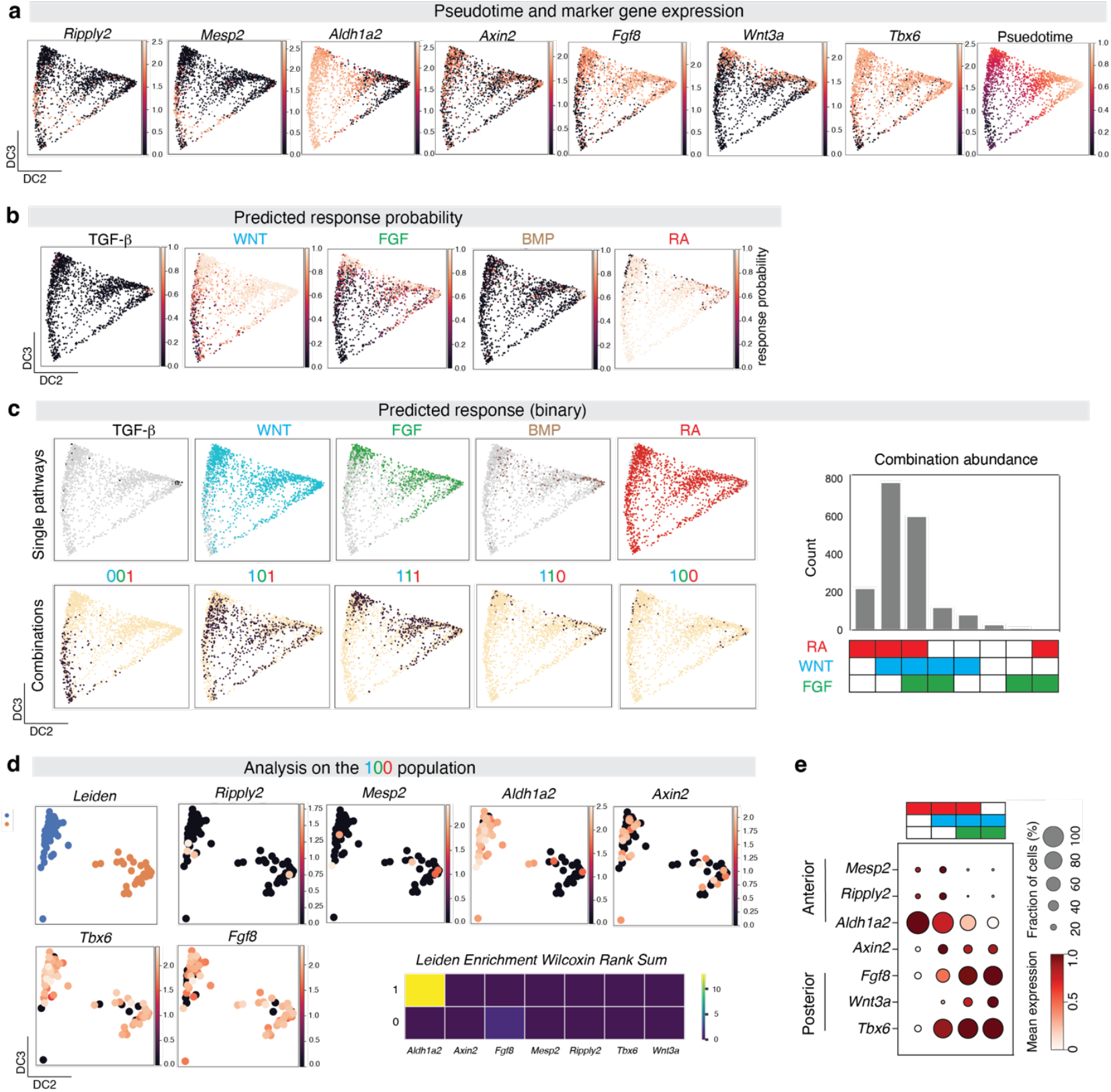
Spatial reconstruction of heterogeneous signaling states among somitic mesoderm cells. **a**, Expression of marker genes that are associated with spatial positions of the cells in the somitic mesoderm (n=2079) plotted on a diffusion map (Anterior: *Ripply2, Mesp2, Aldh1a2*; Posterior: *Tbx6, Wnt3a, Fgf8*; Signaling State: *Axin2*). **b**, Predicted response probability for each signaling pathway on the somitic mesoderm population annotated in the atlas. **c**, Binary predictions of pathway activation and their combinations (*left*). Counts of cells annotated with each signal combination, with top five combinations accounting for 99.5% of the cells (*right*). **d**, Re-clustering of the predicted WNT+FGF-RA-cells show two distinct clusters with either anterior or posterior identities, suggesting WNT+FGF-RA-cells are most likely a collection of cells with false-negative and/or false-positive annotation of the signaling states. **e**, Differential enrichment of marker genes associated with spatial positions among inferred subpopulations of the somitic mesoderm. p<0.05 for RA+WNT-FGF- (*Ripply2, Mesp2, Aldh1a2)*, RA+WNT+FGF*-* (*Aldh1a2, Tbx6)*, RA+WNT+FGF+ (*Wnt3a, Fgf8, Tbx6)* and RA-WNT+FGF+ (*Wnt3a, Fgf8, Tbx6*) (one-tailed Mann-Whitney U-test (alternative: greater) of each gene’s expression in each subpopulation relative to all other subpopulations).

**Extended Data Figure 18.**
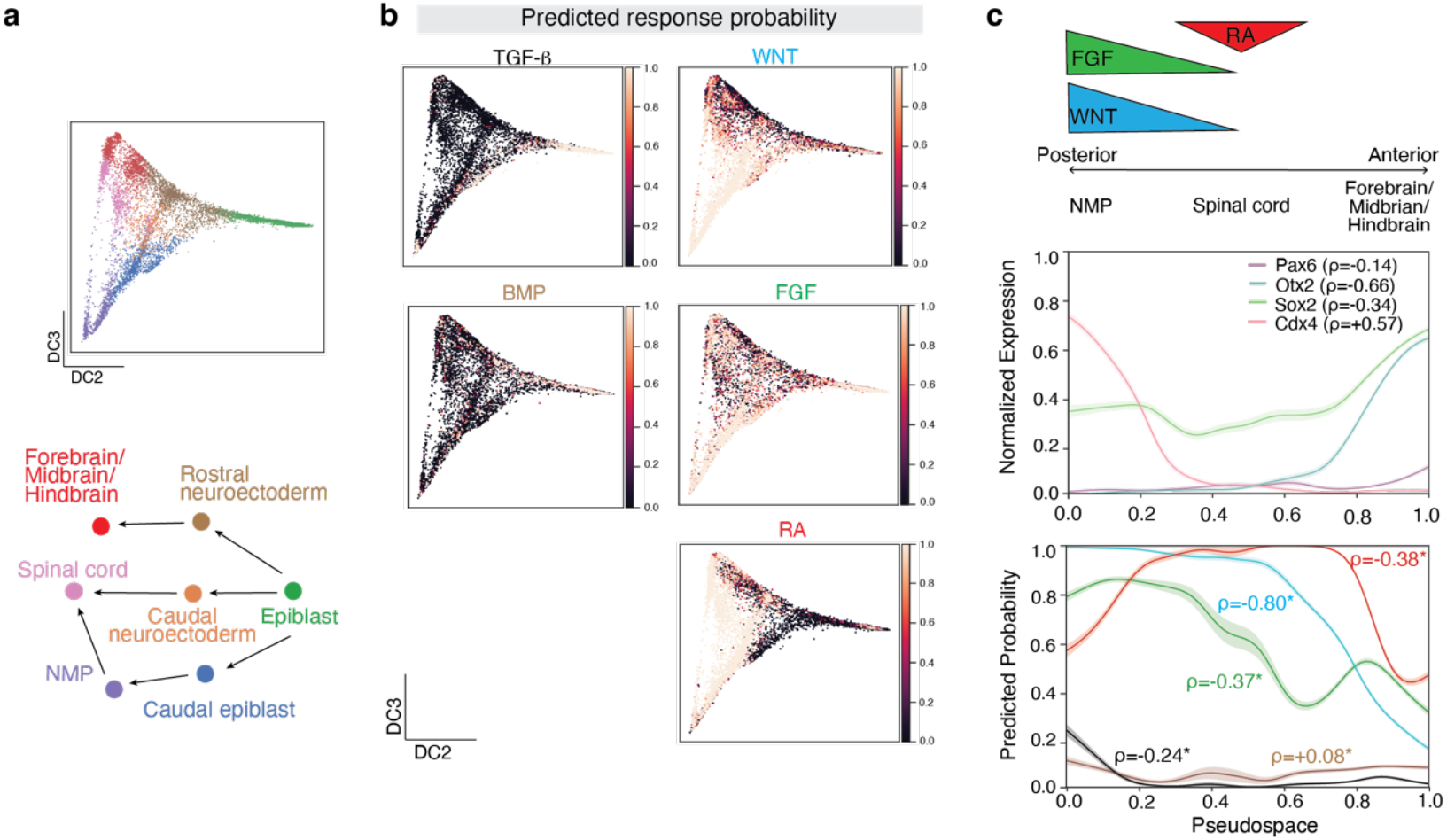
IRIS predictions in early neural differentiation. **a**, Diffusion map of cell populations associated with the early neural lineages (*top*) and a schematic of lineage relationships amongst cell types (*bottom*). **b**, Individual probability predictions for each of the five pathways overlaid on the diffusion map. **c**, Diagram of known spatial gradients during early neural differentiation (*top*). WNT and FGF are known to posteriorize neural fates while BMP and TGF-β family ligands inhibit neural induction in early development ^41,42,56,57^. The role of RA has been well characterized for generation of spinal cord ^41^. NMP (neuromesodermal progenitor)-Spinal cord-Forebrain/Midbrian/Hindbrain cells are aligned along the diffusion pseudospace component, as confirmed by the spatial patterns of marker gene expression (NMP: *Cdx4*; Spinal cord: *Pax6*; Forebrian/Midbrain/Hindbrain: *Otx2, Sox2*) (*middle*). The predicted signaling response probabilities across all five pathways show statically significant spatial trends (*bottom*). * denotes significantly increasing or decreasing trends along spatial axes, assessed by Spearman’s correlation (n=8691, p < 0.05).

**Extended Data Figure 19.**
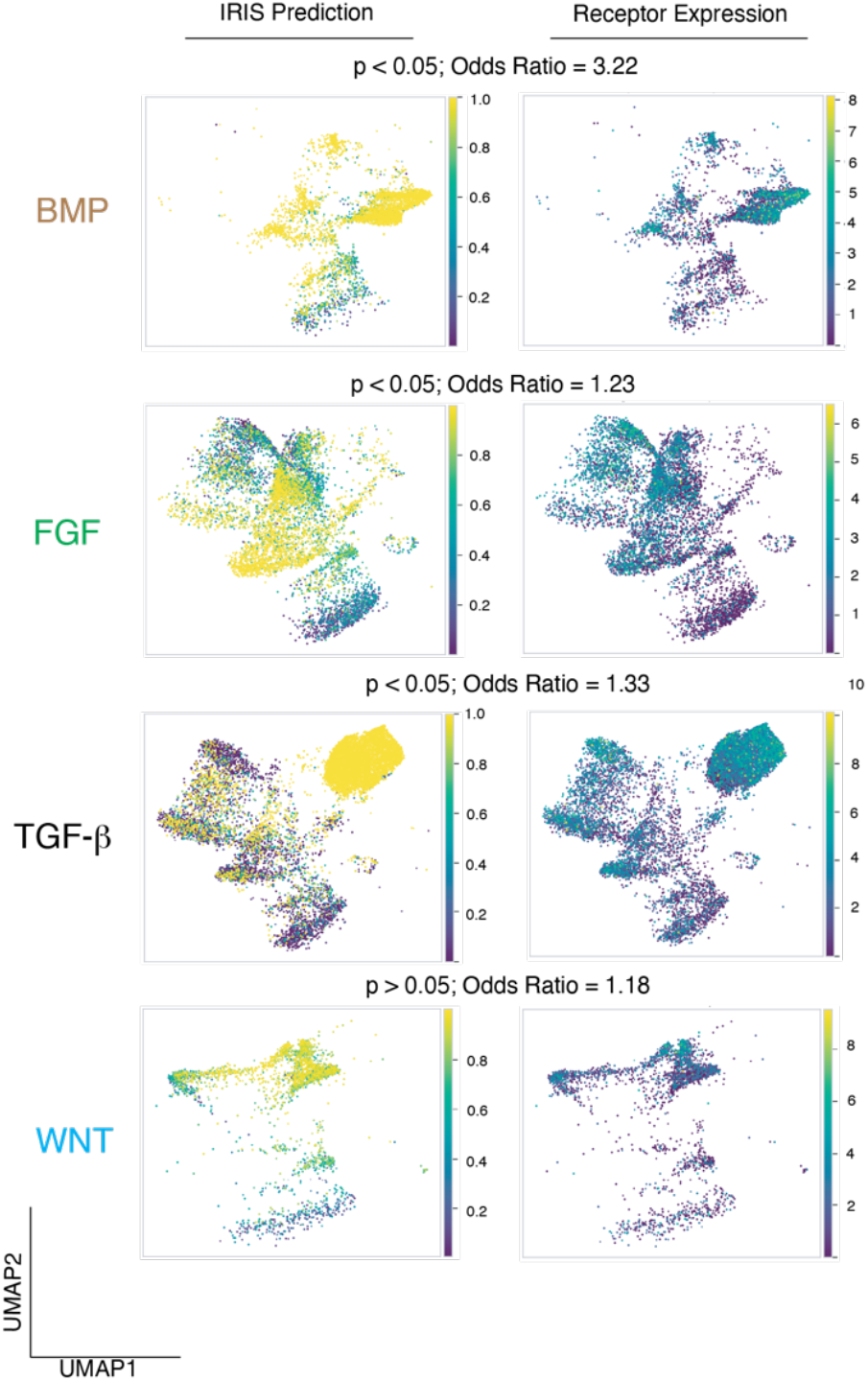
IRIS signaling prediction on adult human airway epithelial cells. IRIS predicted signaling response probabilities on adult human airway epithelial cells, which have been stimulated with the respective ligands for 7 days (*left*). The stimulation includes BMP (BMP4; n=3891), FGF (FGF2 and FGF10; n=7053), TGF-β (TGF-β2 and Activin A; n=11084), and WNT (chemical agonist CHIR; n=2643) ^43^. Summed expression levels of the corresponding receptor families for each pathway in counts-per-million (CPM) (*right*). Cells predicted to be activated by the signaling pathway were significantly enriched among cells with detectable levels of the corresponding receptor expression (p < 0.05; Fischer’s exact test) except for cells treated with CHIR, which bypasses receptors to directly activate the intracellular signal transduction of the WNT pathway. The odds ratio shows the strength of a statistically significant relationship. Note that using receptor expression to approximate signaling potential could lead to false negatives due to dropouts of lowly expressed receptors.

**Extended Data Figure 20.**
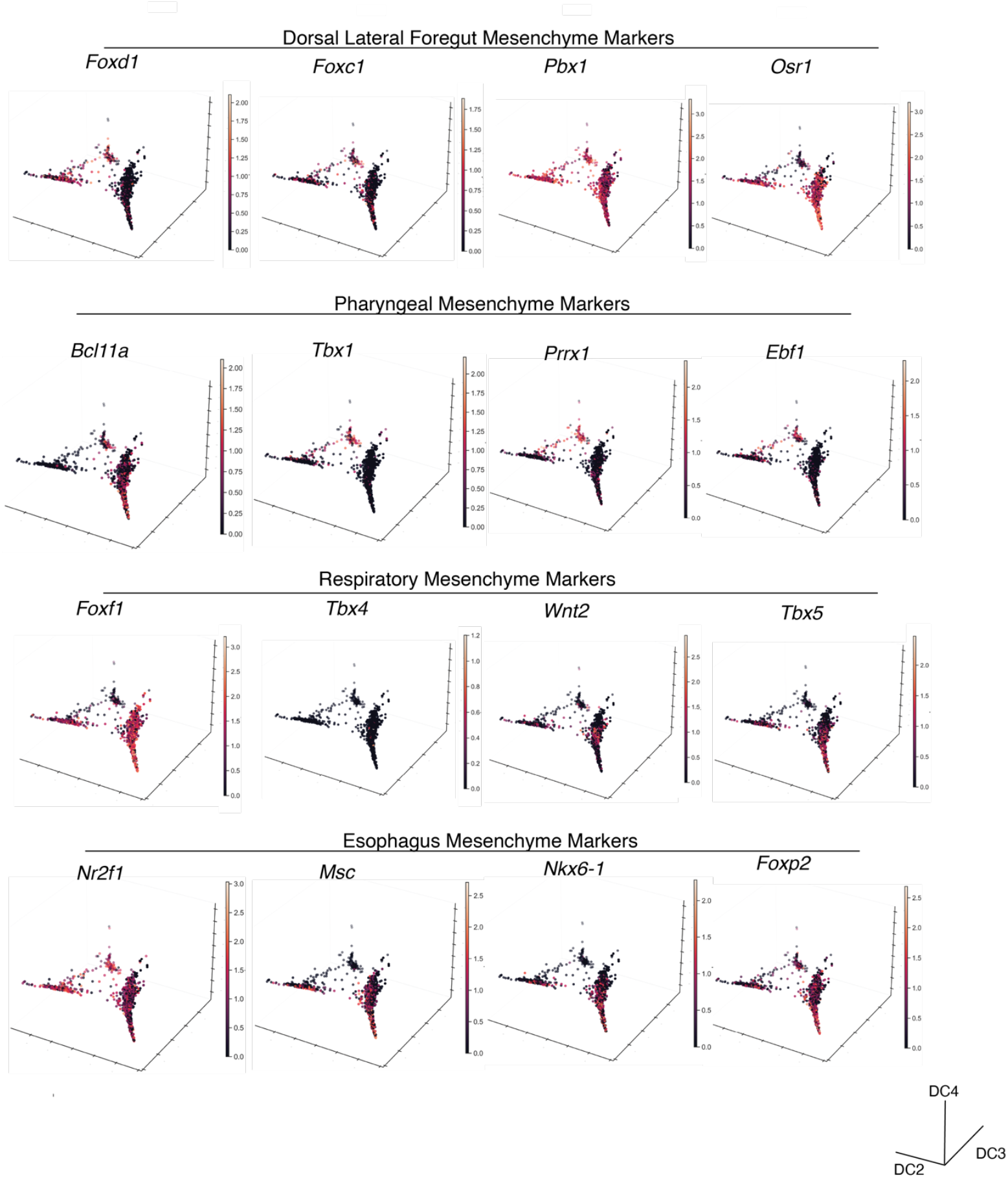
Fate divergence of organ-specific mesenchyme. Marker genes corresponding to the dorsal lateral foregut mesenchyme, esophageal, pharyngeal and respiratory mesenchyme are overlaid on the diffusion maps. The respective cell populations are from E9-E9.5 mouse embryo foregut ^17^, and the 3D diffusion map was constructed in **Fig. 4a**.

**Extended Data Figure 21.**
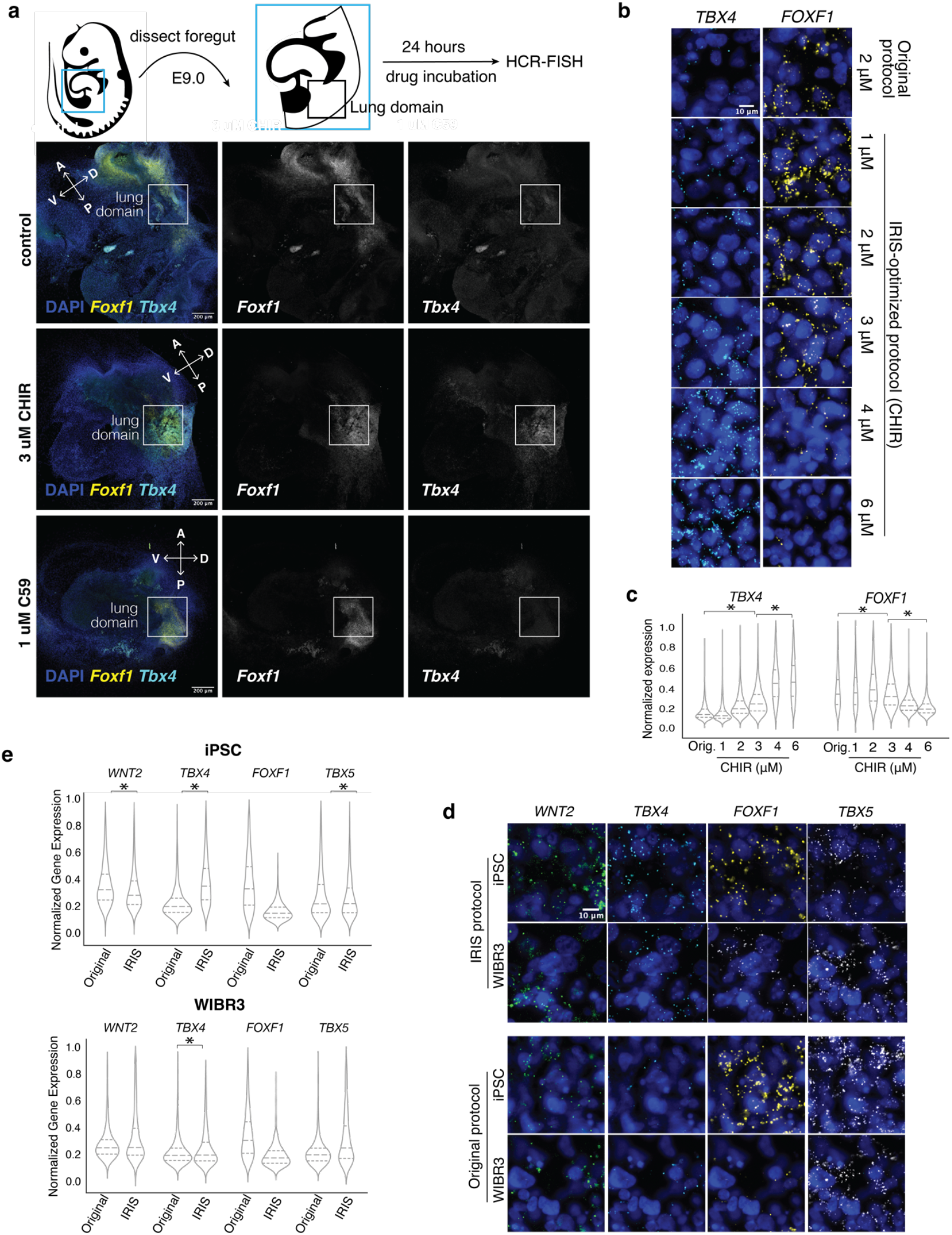
Validation of IRIS prediction on the role of WNT in respiratory mesenchyme specification. **a**, Whole mouse foregut explant images revealed that the expansion of *Tbx4+* domain upon WNT stimulation was restricted to the tissues adjacent to the putative lung domain. Zoomed out images of **Figure 4b** Activation of WNT by CHIR induces an expansion of the *Tbx4+* domain, while treatment of a WNT inhibitor C59 completely ablated the *Tbx4+* domain. **b, c**, Titrating the level of WNT activation in the IRIS-optimized hESC differentiation protocol (WiCells WA01) modulated the expression levels of key respiratory mesenchyme markers. Multiplexed HCR-FISH was performed on Day 7 with representative images shown in (**b**) and quantification of transcripts per cell in (**c**). Number of cells for each condition: 6 μM, n=3228; 4 μM, n=2986; 3 μM, n=2990; 2 μM, n=3040; 1 μM, n=3417; Original, n=3913). **d, e**, IRIS-optimized protocol increased the expression of lung mesenchyme markers at the single-cell level, a result replicated across multiple human pluripotent cell lines. Multiplexed HCR-FISH was performed on Day 7 with representative images shown in (**d**) and quantification of gene expression levels per cell in (**e**). Number of cells for each condition: iPSC with IRIS protocol, n=2421; iPSC with original protocol, n=3200; WIBR3 with IRIS protocol, n=1637; WIBR3 with original protocol, n=2843). * denotes significant p value of <0.05 according to Student’s T-test with the alternative hypothesis of a greater mean.

## Notes

### Competing Interest Statement

The authors have declared no competing interest.

### Summary of Updates

Figures 2 and 3 revised; additional data and analysis included; supplemental figures updated.

